# EV-Elute: a universal platform for enrichment of functional surface marker-defined extracellular vesicle subpopulations

**DOI:** 10.1101/2023.10.25.563755

**Authors:** Willemijn S de Voogt, Rowan Frunt, Raul M Leandro, Casper S Triesscheijn, Bella Monica, Ioanna Paspali, Mark Tielemans, Jerney JJM Francois, Cor W Seinen, Olivier G de Jong, Sander AA Kooijmans

## Abstract

Intercellular communication via extracellular vesicles (EVs) has been identified as a vital component of a steadily expanding number of physiological and pathological processes. To accommodate these roles, EVs are equipped with specific proteins, lipids, and RNA molecules by EV-secreting cells. Consequently, EVs have highly heterogeneous molecular compositions. Given that surface molecules on EVs determine their interactions with their environment, it is conceivable that EV functionality differs between subpopulations with varying surface compositions. However, it has been technically challenging to examine such functional heterogeneity due to a lack of non-destructive methods to separate EV subpopulations based on their surface markers. Here, we used Design-of-Experiments methodology to rapidly optimize a protocol, which we name ‘EV-Elute’, to elute intact EVs from commercially available Protein G-coated magnetic beads. We captured EVs from various cell types on these beads using antibodies against CD9, CD63, CD81 and a custom-made protein binding phosphatidylserine (PS). When applying EV-Elute, over 70% of bound EVs could be recovered from the beads in a pH– and incubation time-dependent fashion. EV subpopulations were found to be devoid of co-isolated protein contaminants observed in whole EV isolates and showed intact morphology by electron microscopy. Proteinase K protection assays showed a mild and reversible decrease of EV membrane integrity during elution. Depending on the type of capturing antibody used, some antibodies remained EV-associated after elution. EV subpopulations showed uptake patterns similar to whole EV isolates in co-cultures of peripheral blood mononuclear cells and endothelial cells. However, in Cas9/sgRNA delivery assays, CD63^+^ EVs showed a lower capacity to functionally deliver cargo as compared to CD9^+^, CD81^+^ and PS^+^ EVs. Taken together, we developed a novel, easy-to-use platform to isolate and functionally compare surface marker-defined EV subpopulations. Importantly, this platform does not require specialized equipment or reagents and is universally applicable to any capturing antibody and EV source. Hence, EV-Elute can open new opportunities to study EV functionality at the subpopulation level.

## Introduction

Extracellular vesicles (EVs) are submicron-sized membrane-enclosed particles secreted by many, if not all cells in the body. Initially EVs were regarded solely as carriers of cellular waste products, but the rapidly expanding EV research field has now implicated EVs in a vast array of physiological and pathological processes. The functionality of EVs in these processes is rather diverse. For example, stem cell-derived EVs are often heralded for their capacity to stimulate cell proliferation in regenerative processes[1, 2], but also growth inhibitory effects of such EVs have been reported[3–5]. In a similar fashion, antigen presenting cell-derived EVs have been shown to aid in the induction of immune responses against pathogens[6, 7], while other studies highlight potent immunosuppressive effects of such EVs[8].

These seemingly opposing effects of EVs can stem from differences in EV source, isolation method and study protocols, however it is also increasingly recognized that EVs isolated from a single cell source consist of a highly heterogenous mixture of EV subpopulations, which potentially each have different biological functions[9]. For example, dendritic cells have been shown to release EV subtypes which induce different T-cell responses[10]. This functional heterogeneity results partly from the intracellular origin of the EVs. Some EVs (referred to as exosomes) are formed by inward budding of the membrane of multivesicular endosomes (MVEs), and are released by fusion of the MVEs with the plasma membrane. This process puts size constraints on exosomes, which typically do not exceed diameters of 200 nm. On the other hand, EVs can be formed via direct budding and fission from the plasma membrane. These EVs are often referred to as microvesicles or ectosomes, and can span a wider size range (50-1000 nm). Finally, apoptotic bodies can be considered the largest class of EVs, often exceeding sizes of 1000 nm and originating from blebbing of the membranes of apoptotic cells[11]. Microvesicles and exosomes are the most studied classes of EVs, but due to their largely overlapping size and molecular composition, they are difficult to fully separate and are therefore often collectively referred to as EVs[12].

EV-secreting cells load EVs with a rich mixture of biologically active components, including various RNA species, soluble, structural and membrane proteins and lipids[13, 14]. The molecular composition of EVs varies within a given size range based on the type and state of the secreting cell, their intracellular origin and potentially their designated function, adding an extra layer of complexity to EV research. As we reviewed elsewhere[15], it is conceivable that the biological function of EVs correlates with the molecules on their surface. EV surface molecules dictate interactions with their environment, and could for example promote immune evasion [16] or clearance[17], cell-specific docking and uptake[18], processing[19, 20] and potentially cargo release [21]. As a consequence, EV isolates derived from gold standard size-based isolation procedures (e.g. ultracentrifugation or size-exclusion chromatography) contain EV subtypes with distinct and/or overlapping surface signatures and diverging functionality. This functional heterogeneity is a formidable challenge for the application of EVs as drug delivery vehicles and significantly complicates studies on EV biology and functionality[22], where EVs are often studied in bulk.

To unravel and compare EV functionality at the level of surface marker-defined subpopulations, it is required to specifically isolate such subpopulations without compromising EV integrity. However, to our knowledge, non-destructive and universally applicable methods for the separation of EV subtypes based on their surface molecules are still lacking. Previous studies have mostly focused on EV subtypes characterized by certain size ranges[23–26] or densities[27–29], as these subtypes are relatively straightforward to separate intactly using routine EV isolation methodology and to perform functional comparisons[10, 30]. However, as discussed above, within specific size ranges EVs still contain heterogeneous surface compositions. Immunoprecipitation is a commonly used technique to further enrich and phenotype EV subtypes containing specific surface markers[26, 31–33]. Unfortunately however, the high-affinity binding of antibodies with their target proteins prohibits the use of immunocaptured EVs for functional studies. Instead, EVs are often stripped from the antibody-coated surface with stringent lysis buffers or low/high pH solutions in order to analyze protein, RNA or lipid content[34–38]. Alternative methods for elution of intact EVs have been proposed, which rely on the use of antibodies modified with ‘releasable’ moieties such as chemical linkers[39], DNA linkers[36] and/or biotin groups[40, 41]. Unfortunately, such strategies have limitations, as modified antibodies against uncommon epitopes are often not commercially available. In addition, custom antibody modification typically requires expensive reagents and may affect antibody functionality.

To overcome these restrictions, we describe a novel platform to (I) isolate EV subpopulations by their surface markers (II) without compromising EV functionality and (III) with universal compatibility with different capturing antibodies and EV sources. To this end, we applied a previously described immunoprecipitation method[26] to capture pre-isolated EVs from three cell lines on commercially available Protein G-coated magnetic beads using antibodies against canonical EV surface proteins CD9, CD63 and CD81. In addition, we used a recombinant Myc-tagged phosphatidylserine (PS)-binding protein which we previously developed[42] combined with anti-Myc antibodies to capture PS^+^ EVs. Then, we tested a library of elution buffers formulated according to Design-of-Experiments (DoE) methodology to identify critical factors for EV elution and EV damage. Through this strategy, we systematically optimized a protocol to effectively elute EVs while minimizing EV damage, which we name ‘EV-Elute’. We used EV-Elute to study the cell-specificity and uptake of the abovementioned EV subpopulations in co-cultures of primary immune cells and endothelial cells. Finally, we compared their capacity to functionally deliver engineered therapeutic cargo (Cas9/sgRNA ribonucleoprotein complexes) to recipient cells.

## Materials and methods

### Molecular cloning

Expression cassettes for FLAG-tagged CD9 or CD63 fused to the MS2 bacteriophage coat protein (MCP) via a photocleavable linker (CD9– or CD63-PhoCl-MCP, respectively) were cloned into a pHAGE2 expression vector containing a puromycin resistance gene as described elsewhere [Elsharkasy et al., manuscript in preparation]. Similarly, cloning of a plasmid driving expression of a sgRNA modified with MS2 aptamers in the second stemloop and tetraloop from a U6 promoter is described previously[43]. Plasmids encoding the spike glycoprotein of vesicular stomatitis virus (VSV-G) and Cas9 were obtained from Addgene (cat. 12259 and cat. 52962, respectively). A pHAGE2 vector containing a fluorescent stoplight expression cassette (pHAGE2-Stoplight) which is targetable by Cas9 was generated as previously described[44].

### Cell culture

All used cell lines were maintained at 37°C and 5% CO_2_. HEK239T, HeLa and MDA-MB-231 cell lines were purchased from the American Type Culture Collection (ATTC) and cultured in Dulbecco’s Modified Eagle Medium with L-glutamine (DMEM, Gibco) supplemented with 10% fetal bovine serum (FBS) and 1% penicillin/streptomycin (Gibco). Human endothelial cells (HMEC-1) stably expressing enhanced green fluorescent protein (eGFP) were generated as described before[45] and cultured on 0.1% gelatin (VWR) pre-coated surfaces in MCDB-131 medium (Gibco) supplemented with 10% FBS, 1% penicillin/streptomycin, 10 ng/mL human epidermal growth factor (hEGF, PreproTech), 50 nM hydrocortisone (Sigma Aldrich), 2 mM L-glutamine (Gibco) and 0.5 mg/mL G418. HEK293T cells stably lentivirally transduced with pHAGE-Stoplight (HEK293T-Stoplight) were generated as described before[44] and were cultured in DMEM supplemented with 10% FBS, 1% penicillin/streptomycin and 1 mg/mL G418. To generate EVs loaded with Cas9 and sgRNA, HEK293T cells were grown to a confluency of 80% and transfected with a mixture of plasmids encoding CD9– or CD63-PhoCl-MCP, Cas9, MS2 aptamer-modified sgRNA and VSV-G in a ratio of 2:2:2:1 at a dose of 20 µg total plasmid DNA using 50 µg linear 25K MW polyethylenimine (PEI, Polysciences) per T175 flask. After 24 hours, cells were washed and EV production and isolation protocols were initiated as described below.

### EV isolation

For EV production, HEK293T, HeLa or MDA-MB-231 cells were cultured in T175 flasks until a confluency of 70-80% was reached. Cells were washed once with phosphate buffered saline (PBS, Sigma) and cultured in OptiMEM with GlutaMAX (Gibco) supplemented with 1% penicillin/streptomycin for 16-20 hours. Medium was collected in 50 mL tubes and centrifuged for 10 min at 300 x *g* at 4°C. Supernatants were centrifuged for 10 min at 2000 x *g* at 4°C, and resulting supernatants were filtered through 0.45 µm aPES Rapid-Flow bottle top filters (Nalgene). Medium was kept on ice and concentrated 20-40 times to a volume of 15-20 mL by tangential flow filtration (TFF) using Minimate TFF capsules with 100 kDa molecular weight cut-off (MWCO) Omega membranes coupled to a Minimate EVO TFF system (kindly provided by Pall Life Sciences). Medium was further concentrated to volumes <1 mL using 100kDa MWCO Amicon Ultra-15 Centrifugal Filter Units (Merck Millipore). Tricorn 10/300 chromatography columns (Cytiva) packed with Sepharose 4 Fast Flow resin (Cytiva, column volume ±24mL) according to manufacturer’s instructions were equilibrated with PBS at 1 mL/min using a refrigerated ÄKTA start or ÄKTA Pure chromatography system. Concentrated medium samples were injected and 1 mL fractions were collected. EV-containing fractions (identified by UV absorbance at 280 nm, typically fractions 8-12) were pooled, filtered through 0.45 µm syringe filters (for Cas9/sgRNA delivery experiments) and concentrated using 100kDa MWCO Amicon Ultra-4 Centrifugal Filter Units (Merck Millipore) to volumes of 100-250 µL.

### EV fluorescent labelling

For cell uptake and Cas9 delivery assays, EVs were labelled with MemGlow 590 Fluorogenic probe, whereas for single EV imaging, EVs were labelled with Memglow 560 Fluorogenic probe (Cytoskeleton, Inc). MemGlow was dissolved as 20 µM stock solutions in dimethyl sulfoxide (DMSO) and diluted to 400 nM in PBS. Labelling solution was mixed in a 1:1 ratio with EVs and incubated for 30 min at room temperature. Subsequently EVs were loaded onto a PBS pre-equilibrated XK-16/20 column (Cytiva) which was packed with Sepharose CL-4B resin (Merck) according to manufacturer’s instructions. EV-containing fractions were collected using an ÄKTA chromatography system and concentrated as described above.

### Nanoparticle Tracking Analysis (NTA)

EVs were diluted to appropriate dilutions (40-100 particles/frame) using PBS and injected into a NanoSight NS500 system equipped with an LM14 405 nm laser unit (Malvern Instruments). Five 30-second movies were recorded at camera level 15 at a fixed temperature of 25°C and analyzed with NTA 3.4 software at detection threshold 7. Measured particle concentrations were corrected for particle concentrations measured in PBS used for sample dilution.

### EV capture on magnetic beads and elution

EVs were incubated with antibodies against CD9, CD63 or CD81 (Supplementary Table 1) in fixed, titrated ratios for 30 min at room temperature: 1 x 10^10^ particles as measured by NTA were mixed with either 30 ng anti-CD9, 300 ng anti-CD63, or 40 ng anti-CD81 antibodies. Alternatively, 1 x 10^10^ EV particles were incubated with 170 ng of previously generated, Myc-tagged non-binding nanobodies fused to PS-binding C1C2 domains of mouse lactadherin[42] (R2-C1C2) for 15 min, followed by incubation with 200 ng anti-Myc antibody (Supplementary Table 1) for 15 min at room temperature. Dynabeads Protein G (Thermo Fisher Scientific, 1.67 µL per 1 x 10^10^ EV particles) were washed three times with PBS containing 0.001% v/v Tween20 (PBS-T) and once with PBS using a magnetic rack.

Supernatant was removed and beads were incubated with EV/antibody mixtures in total volumes of 40-80 µL overnight at 4°C on a rotator. The next day, beads were placed on a magnet and supernatants were collected. Beads were washed five times with PBS-T and supernatant was removed. For EV elution, beads were incubated for indicated times in 50-100 µL of selected triethylamine (TEA)-based elution buffers (Supplementary Table 2) or EV-Elute buffer (145.4 mM sodium chloride (NaCl), 50 mM disodium hydrogen phosphate (Na_2_HPO_4_), 12.5 mM sodium hydroxide (NaOH), pH 11.2). Eluted EVs were immediately mixed with 0.15 volume neutralization buffer (1M Tris hydrochloride (Tris-HCl), pH 4.1) on ice. For capture experiments where bead-bound EVs were analyzed by flow cytometry, 6-fold lower volumes of Dynabeads Protein G were used to capture EV/antibody mixtures described above. In Cas9/sgRNA delivery assays, aliquots of EV/antibody mixes were incubated overnight separately in the absence of magnetic beads as controls. The next day, controls were treated with 10 volumes of EV-Elute buffer for 1 min, mixed with 0.15 volumes of neutralization buffer and further processed the same as eluted EV subpopulations.

### DoE-based generation of library of elution conditions

A DoE-based library of elution conditions was generated using the ‘Custom Design’ module of JMP 17.1.0 software (SAS). TEA concentration, incubation time and pH were selected as continuous factors with ranges of 0.5 – 20 mM, 5 – 30 min and pH 10 – 11.5, respectively. For MDA-MB-231 EVs, a pH range of 9 – 11.5 was adopted. ‘Elution efficiency’ and ‘EV permeability’ were selected as response parameters with 0 – 100% ranges and goals to maximize and minimize, respectively. The response surface methodology (RSM) function was employed to test both main effects and interactions between the three factors. The resulting design consisted of 15 unique elution conditions (Supplementary Table 3). Elution buffers were formulated by diluting TEA to appropriate concentrations with PBS and adjusting pH using sodium hydroxide (NaOH) and/or hydrochloric acid (HCl). Elution efficiency was tested in triplicate using flow cytometry and EV permeability was assessed by western blotting in duplicate (see below). Experimental data on EV elution and permeability was expressed as a percentage of control conditions. Using JMP software, for every response parameter a model was fitted using the standard least squares personality with emphasis on effect screening. Non-significant effects were removed from the model and resulting models were used to generate prediction formulas for each response parameter.

### Analysis of beads using flow cytometry

EVs captured on Dynabeads Protein G were washed and/or eluted as described above. EV-coated beads ‘mock eluted’ with PBS were included as controls. Beads were transferred to a U-bottom 96-well plate (Greiner) and pulled down on a Magnetic Plate Separator (Luminex). Beads were resuspended in 50 µL Diluent C mixed with 1:550 diluted PKH67 (Sigma Aldrich) per well, and incubated for 5 min at room temperature shielded from light. Labelling reactions were quenched by addition of 150 µL 1% bovine serum albumin (BSA) in PBS (1% PBS-A). Supernatant was removed, beads were washed once with 1% PBS-A, once with PBS-T and finally resuspended in PBS-T. Twenty thousand beads per well were acquired using a FACSCanto II flow cytometer (BD Biosciences) Data was analyzed using FlowJo v10.9.0. Single beads were gated according to gating strategy in Supplementary Figure 1 and mean fluorescence intensity (MFI) of the PKH67 dye was determined. The MFI of empty beads was measured as blank and subtracted from samples to calculate ΔMFI. Percentage of EV elution was calculated using the following formula:

Elution efficiency (%) = 100 – (100 x ΔMFI_Sample_ / ΔMFI_PBS mock elution_)

### Proteinase K protection assay and western blotting

EVs (± 1 x 10^10^ particles per sample) were diluted with 10 volumes of selected elution buffers for indicated durations in the presence of 20 µg/mL Proteinase K (Sigma) at 37°C. Reactions were stopped by addition of 5 mM phenylmethanesulfonyl fluoride (PMSF, Sigma, dissolved as 140 mM stock solution in isopropanol) and 0.15 volumes of neutralization buffer (1M Tris-HCL). As a positive control for full intraluminal protein digestion (100% EV permeability), EVs were mixed with proteinase K and 0.5% sodium dodecyl sulfate (SDS) in PBS. EVs mixed with PBS and proteinase K served as negative controls (0% EV permeability). Samples were mixed with reducing sample buffer containing dithiothreitol (DTT) if needed, incubated at 85°C for 10 min and loaded on Bolt 4-12% Bis-Tris polyacrylamide gels (Thermo Fisher Scientific). Proteins were electrotransferred to Immobilon-FL polyvinylidene difluoride (PVDF) membranes (Merck Millipore) and blocked with Intercept Blocking Buffer (LI-COR Biosciences) mixed in a 1:1 ratio with Tris buffered saline (TBS). Membranes were stained with primary and secondary antibodies (Supplementary Table 1) diluted in Intercept Blocking Buffer diluted 1:1 with TBS containing 0.1% Tween20 (TBS-T). Blots were imaged using an Odyssey M imager (LI-COR Biosciences) at 700nm and 800nm channels. Band intensities were quantified using Empiria Studio version 2.1.1.138 (LI-COR Biosciences). In order to assess EV permeability, the band intensities of intraluminal EV proteins Alix, TSG-101, syntenin and β-actin were compared with the band intensities of these proteins in negative control samples (EVs incubated with proteinase K in PBS, for the same duration) on the same membrane. Percentage of EV permeability was calculated for each individual intraluminal protein using the following formula:

EV permeability (%) = 100 – (100 x Band intensity_Sample_ / Band intensity_Negative control_)

Average EV permeability values obtained from bands of Alix, TSG-101, syntenin and β-actin were calculated and used as input for model fitting in JMP (see above).

### Immunogold labelling and electron microscopy

Ten microliters of EVs were adsorbed to carbon-coated formvar grids for 20 min at room temperature. Grids were washed with PBS and blocked with blocking buffer (0.5% fish skin gelatin (Sigma) and 0.1% BSA-c (Aurion) in PBS) for immunogold labelling, or fixed using 1% glutaraldehyde in PBS. Immunogold labelling was performed by staining with rabbit-anti-mouse IgG (Rockland, cat. 610-4120, 1:250) followed by 10 nm Protein A-gold (CMC, Utrecht, The Netherlands) and fixation using 1% glutaraldehyde in PBS. After excessive washing with MilliQ water, grids were counterstained with uranyl-oxalate and embedded in methyl cellulose uranyl-acetate. Imaging was performed using a Tecnai T12 transmission electron microscope (FEI). For gold quantification, 5-10 pictures were acquired at random grid locations at a magnification of 49.000x. A custom-made ImageJ script based on intensity thresholding was run on each picture to count the total number of gold particles.

Membrane-associated gold was counted manually using the multi-point tool in ImageJ and expressed as a percentage of total gold.

### Cryogenic electron microscopy

Ten microliters of EVs were incubated on freshly glow-discharged C-flat Holey carbon grids (Electron Microscopy Sciences) for at least 10 min in a humid chamber. Vitrification was performed using a Mark IV Vitrobot (FEI) and samples were stored in liquid nitrogen until imaging. Grids were loaded in a liquid nitrogen-precooled 70° tilt cryo-transfer system (Gatan, Inc.) and inserted in a Tecnai G2 20 TWIN 200kV transmission electron microscope (FEI). Images were acquired using a bottom-mounted High-Sensitive 4k x 4k Eagle CCD Camera System (FEI) at 200 kV.

### Direct stochastic optical reconstruction microscopy (dSTORM) imaging

Antibodies against CD9, CD63, CD81 and IgG control antibodies (Supplementary Table 1) were depleted from sodium azide using a 10kDa MWCO filter ultrafiltration vial (Biotium). Ten microgram of each antibody was conjugated overnight at 4°C with 40µg/mL NHS-AlexaFluor647 (AF647, ThemoFisher Scientific) in 0.1M borate buffer (pH 8.6). Subsequently, free NHS-AF647 was eliminated with 0.5mL 7kDa MWCO Zeba Dye and Biotin Removal Spin Columns (Thermo Fisher Scientific). Conjugated antibodies were used immediately or stored at 4°C. Coverslips (#1.5H, Marienfeld) were prepared for dSTORM imaging by ultrasonication for two times 30 min in MilliQ water and 30min in 1M potassium hydroxide. After drying the coverslips with nitrogen gas, 50-well CultureWell^TM^ gasket (Grace Bio-Labs) were attached creating 10µL wells. Coverslips were coated with 0.01% poly-L-lysine (PLL) at 37°C for 2h. Memglow 560-labelled EVs were incubated on the coverslips overnight at 4°C. The next day, coverslips were blocked in 5% PBS-A for 1h at room temperature and stained with 10µg/mL of AF647-conjugated antibodies O/N at 4°C. Coverslips were washed twice with PBS and imaged on the same day. dSTORM imaging was performed using a Nanoimager-S (ONI) equipped with 405nm (150 mW), 488nm (1W), 561nm (500 mW), 640nm (1W) lasers and a 100x objective with numerical aperture 1.4 (Olympus). Prior to acquisition, channel mapping was performed with a microscopy slide containing 0.1µm TetraSpeck^TM^ beads (Thermo Fisher Scientific). Tetraspanin dSTORM imaging was performed in freshly prepared OxEA buffer[46] with pH 8-8.5. In brief, a TIRF angle was applied and 7500 frames were acquired at 640nm (laser power 35%) with 30ms exposure time. Subsequently 500 frames were acquired at 561nm (laser power 30%) with 30ms exposure time to image the EV reference MemGlow 560 signal. Filtering of localizations and HDBSCAN cluster analysis was performed in the collaborative discovery (CODI) online analysis platform (https://alto.codi.bio/, ONI). For both fluorescent channels, the following filters were applied: Drift correction: DME; photon count: > 300; localization precision: 0-20 nm; p-value: >1 x 10^-6^. The following HDBSCAN clustering parameters were used: area: < 80.000 nm^2^, skew: < 2.6; kappa: >23 nm; # of localizations: > 14. One hundred twenty nanometer radius centroids were applied for counting localizations in individual clusters whereby > 2 localizations in the 561nm channel and > 12 localizations in the 640nm channel were considered positive. The number of AF647^+^/Memglow 560^+^ clusters was expressed as a percentage of all Memglow 560^+^ clusters.

### EV uptake assays

HMEC-1-eGFP cells were seeded at 15.000 cells/well in a gelatin-coated 96-well plate. The next day, EDTA-anticoagulated blood was obtained from healthy volunteers. Blood withdrawal was approved by the University Medical Center Utrecht Ethics Committee. Informed consent was obtained from all healthy volunteers. Blood was transferred to Ficol-Paque PLUS (GE Healthcare)-loaded Sepmate-50 tubes (Stemcell Technologies) according to manufacturer’s instructions and centrifuged at 12.000g for 10 min at room temperature. Peripheral blood mononuclear cell (PBMC)-containing fractions were collected in wash buffer (2% FBS in PBS) and washed twice at 300 x *g* for 8 min at room temperature. PBMCs were suspended in RPMI 1640 (Gibco) supplemented with 10% FBS and 1% penicillin/streptomycin. Culture medium was removed from HMEC-1-eGFP cells and PMBCs were seeded in the same wells at a density of 300.000 cells/well. Concentrations of EVs and EV subpopulations labelled with Memglow 590 were normalized based on fluorescence measured in a black 384-well plate at an excitation of 573nm and emission of 613nm using a SpectraMax iD3 plate reader (Molecular Devices). EVs and EVs subpopulations were added to the cells and incubated for 4 hours at 37°C. PBMCs were harvested and pooled with PBS-washed and trypsinized HMEC-1-eGFP cells in U-bottom 96-well plates. Cells were washed once with PBS and incubated in 50 µL PBS containing human Fc block (Supplementary Table 1) and Zombie Aqua viability dye (1:200, BioLegend) for 15 min at 4°C. Subsequently 50 µL of antibody cocktail (Supplementary Table 1) in 2% PBS-A was added and incubated for 30 min at 4°C. Cells were washed twice in PBS and fixated with 4% paraformaldehyde in PBS. After washing with PBS, cells were stored overnight in 2% PBS-A. The next day, cells were analyzed using a 4-laser LSRFortessa flow cytometer (BD Biosciences) until data of at least 2.000 cells of each type per well was acquired. Data was analyzed using FlowJo v10.9.0 and an appropriate compensation matrix was applied. Live single cells were classified as follows: HMEC-1: eGFP^+^, CD45^-^; B-cells: CD45^+^, CD19^+^; T-cells: CD45^+^, CD19^-^, CD3^+^; Monocytes: CD45^+^, CD19^-^, CD3^-^, CD14^+^, CD56^-^; natural killer (NK) cells: CD45^+^, CD19^-^, CD3^-^, CD14^-^, CD56^+^; dendritic cells (DCs): CD45^+^, CD19^-^, CD3^-^, CD14^-^, CD56^-^, CD11c^+^. Gating strategy of each cell type is shown in Supplementary Figure 8A. For each cell type, the Memglow 590 MFI of untreated cells was subtracted from the MFI of EV-treated cells to obtain normalized EV uptake ΔMFI data. Cell type-specificity was calculated by expressing the Memglow 590 ΔMFI of each cell type as a percentage of the sum of ΔMFI values over all analyzed cell types.

### Cas9/sgRNA delivery assays

HEK293T cells were transfected to express Cas9, MS2 aptamer-modified sgRNA, CD9 or CD63 fused to MCP via a photocleavable linker, and VSV-G as described above. EVs were isolated, labelled with Memglow 590 and purified as described. HEK293T-Stoplight reporter cells were seeded at a density of 9.000 cells/well in a 96-well plate. The next day, concentrations of EVs and eluted subpopulations were normalized based on Memglow 590 fluorescence, irradiated with 395 nm UV-light using a 50W LED panel (SJLA, China) 20 min on ice, and added to the HEK293T-Stoplight cells. After 4 days, mCherry and eGFP expression was visualized using a 4-channel EVOS M5000 fluorescence microscope (Thermo Fisher Scientific). Cells were harvested by trypsinization, washed two times with FACS buffer (0.3% BSA in PBS) and analyzed using a FACSCanto II flow cytometer (BD Biosciences) until data of at least 70.000 cells per well was acquired. Single mCherry^+^ and eGFP^+^ cells were gated using Flowjo v10.9.0 (gating strategy shown in Supplementary Figure 7B). Reporter activation was expressed as the percentage of mCherry^+^ eGFP^+^ cells of all mCherry^+^ cells.

### sgRNA quantification using RT-qPCR

After fluorescence-assisted concentration normalization, equal volumes of whole EV isolates, EV subpopulations and magnetic bead supernatants were lysed using TRIzol LS Reagent (Invitrogen) and spiked with 25 attomol of a synthetic non-targeting sgRNA (NT sgRNA). RNA was isolated according to manufacturer’s instructions with minor modifications: GlycoBlue Coprecipitant was added to the aqueous phase prior to isopropanol precipitation, and isopropanol precipitation was carried out overnight at –20°C. RNA was washed with 80% ethanol in nuclease-free water, air-dried and dissolved in nuclease-free water. DNAse digestion was performed using Turbo DNA-*free* kit (Invitrogen) according to manufacturer’s instructions to remove potential plasmid contamination. Reverse transcription was performed using SuperScript IV reverse transcriptase (Invitrogen) as follows: 3.5 µL of RNA was mixed with 0.5 µL of 10 mM dNTP Mix (Applied Biosystems), 0.25 µL of reverse primer against stoplight-targeting sgRNA (T-sgRNA), 0.25 µL of reverse primer against NT-sgRNA and 2 µL of nuclease-free water (Ambion). Mixtures were heated to 65°C for 5 min, then incubated on ice for at least 1 min. Then 3.5 µL of reverse transcriptase mix, consisting of 2 µL of 5x RT Buffer, 0.5 µL of 100 mM DTT, 0.5 µL of RNAsin RNAse inhibitor (Promega) and 0.5 µL of Superscript IV reverse transcriptase was added. Reverse transcription was performed for 45 min at 52°C and 10 min at 80°C. cDNA was diluted 5 times with nuclease-free water. For each target gene (T-sgRNA and NT-sgRNA), 3 µL of diluted cDNA was mixed with 0.1 µL of 25 µM forward primer and reverse primer, 5 µL of 2x iQ SYBR Green Supermix (Bio-Rad) and 1.8 µL of nuclease-free water. qPCR was performed in a C1000 Touch thermal cycler equipped with a CFX96 Optical Reaction module (Bio-Rad) with the following cycling program: 3 min at 95°C, followed by 45 cycles of 15 sec at 95°C and 30 sec at 60°C, followed by melt curve analysis. Ct values of T-sgRNA were normalized to those of NT-sgRNA (ΔCt) and converted to fold change compared to whole EV isolates using the 2^-ΔΔCt^ method.

### Statistical analysis

Statistical analysis on data generated from DoE libraries was performed using JMP 17.1.0 as described above. Other statistical analyses were performed using Graphpad Prism 9.3.0.

Comparisons between two groups were made using unpaired two-sided Student’s t tests. Comparisons between more than two groups were made using one-way ANOVA with appropriate posthoc test (see Figure captions). Differences with *p* values < 0.05 were considered statistically significant.

## Results

### DoE-based optimization of EV elution from magnetic beads with minimal EV damage

To screen for a suitable protocol to elute intact EVs from antibody-coated magnetic beads, we isolated EVs from MDA-MB-231 cells using tangential flow filtration (TFF) followed by size-exclusion chromatography (SEC) and captured the EVs on Protein G-coupled magnetic beads using anti-CD9 antibodies. After washing away unbound EVs, we exposed the beads to a range of commonly used buffers in immunoaffinity purification for 10 minutes, stained remaining lipids on the beads with the lipophilic dye PKH67 and analyzed bead fluorescence using flow cytometry (Supplementary Figure 1A). We selected buffers that did not contain organic or denaturing compounds or detergents, as these would be expected to compromise the EV membrane or surface proteins, and ultimately EV functionality. Instead, we focused on buffers that could potentially break the bond between antibody and antigen due to low pH, high pH or high salt concentration. However, our data indicated that neither of these factors predicted efficient EV elution (Supplementary Figure 1B), as buffers with similar pH or salt concentrations resulted in opposite elution patterns. After closer examination of particle size and morphology eluted by the best performing buffers by Nanoparticle Tracking Analysis (NTA) and transmission electron microscopy (TEM), a 10-min exposure to a high pH buffer (pH 11.5) containing 100mM triethylamine (TEA) was selected as the lead elution protocol to take forward to functional studies. However, unfortunately we discovered that this elution protocol caused cell toxicity, prohibiting further use for EV functional studies (data not shown).

To further finetune the elution protocol to be compatible with EV functionality assays and identify factors that are critical for both EV elution efficiency and EV damage, we used Design-of-Experiments (DoE) methodology. This statistical methodology allows exploration of a large range of possible combinations of factors that affect a given process in a time– and cost-effective manner, and predict optimal combinations in a single screening round. In this case, we investigated the effect of three factors in our lead elution protocol on EV elution efficiency and EV damage: incubation time of the buffer with the beads, buffer pH and buffer TEA concentration. For each of these factors, lower and upper boundaries were set based on our previous observations. Using statistical modeling software JMP, a library of 15 elution protocols was generated (Supplementary Table 2 and Figure 1A), which strategically combined these three factors to allow analysis of their individual effect on EV elution, but also interaction effects (e.g. whether a higher buffer pH synergistically acted with longer incubation time to improve EV elution) in a single round of experiments. To test the effect of this library on EV elution, EVs isolated from MDA-MB-231 or HEK293T cells were captured on magnetic beads using antibodies against CD9 or CD81. Beads were exposed to each elution protocol in the library (termed ‘runs’) and the resulting elution was measured using flow cytometry (Figure 1B). In parallel, to assess EV damage, whole EV isolates derived from these cell lines were exposed to each elution protocol in the presence of proteinase K (Figure 1A). We reasoned that, if EV membranes would be perturbed by the elution protocol, proteins in the EV lumen would be degraded by proteinase K. Proteinase K-induced degradation of intraluminal proteins β-actin, Alix, TSG-101, syntenin and caveolin-1 was therefore evaluated using western blotting. Indeed, when EV membranes were compromised by addition of 0.5% SDS, complete degradation of these proteins was observed, even for short incubation times and at harsh conditions (high TEA concentrations at high pH), indicating that proteinase K activity was not inhibited by the elution protocol (Figure 1C and Supplementary Figure 2A). Band intensities of highly abundant intraluminal proteins in each EV type (Alix, β-actin, TSG-101 and syntenin in HEK293T EVs and β-actin, syntenin and caveolin-1 in MDA-MB-231 EVs, Supplementary Figure 2B) were quantified and compared with proteinase K-treated EVs exposed to PBS as controls.

**Figure 1:**
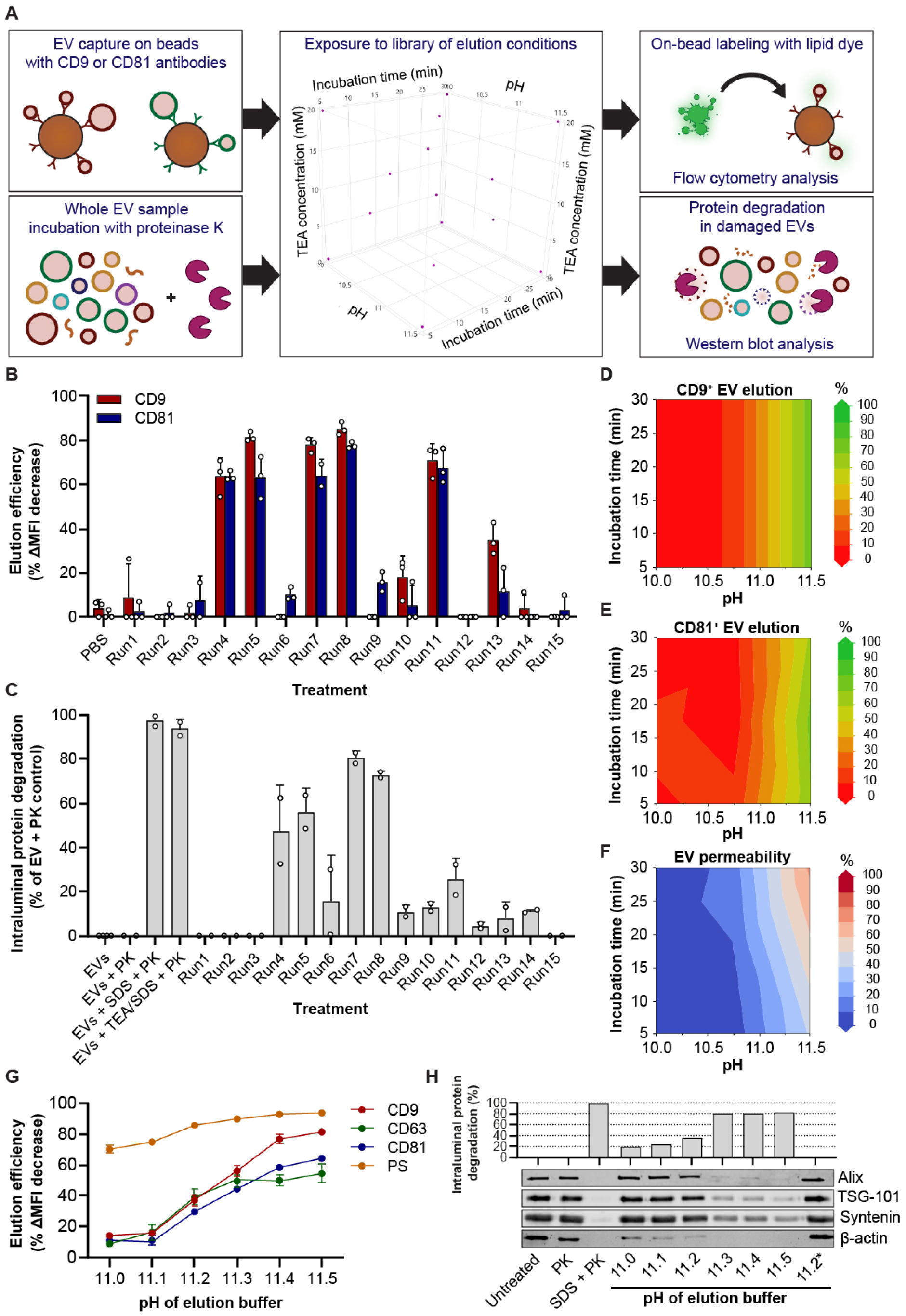
Systematic optimization of EV elution protocol from magnetic beads using DoE methodology. **A:** Overview of experimental setup to test elution of EVs captured on Protein G-coated magnetic beads and EV permeability after exposure to a DoE-based library of elution protocols. Elution protocols varied in incubation time, elution buffer pH and elution buffer TEA concentration. EV elution after each protocol was analyzed by on-bead lipid fluorescent staining and flow cytometry. EV permeability was tested by exposing EVs to proteinase K during each elution protocol and western blotting for intraluminal EV proteins. **B:** Elution efficiency of HEK293T EVs captured using antibodies against CD9 (red) or CD81 (blue) after exposure to PBS as controls or each elution protocol (Run1-15). Mean ± SD of three technical replicates is shown. **C:** HEK293T EV permeability during each elution protocol. Bars show mean ± SD of two technical replicates and indicate average degradation of Alix, β-actin, TSG-101 and syntenin, expressed as a percentage of proteinase K (PK)-treated EVs exposed to PBS. EVs exposed to SDS at physiological pH, or SDS at high pH and TEA concentrations (TEA/SDS) served as positive controls for PK activity. **D, E:** Prediction models of EV elution after capture with anti-CD9 (D) or anti-CD81 (E) antibodies, generated based on data in panel B. **F:** Prediction model of EV permeability generated based on data in panel C. **G:** Elution of HEK293T EVs captured on beads using antibodies against CD9, CD63, CD81 or PS, using elution buffers with pH ranging from 11.0 to 11.5 for 5 min. **H:** Representative western blot of HEK293T EV permeability analysis based on Alix, TSG-101, syntenin and β-actin. EVs were exposed to elution buffers with pH ranging from 11.0 to 11.5 or SDS as control in presence of PK for 5 min. Alternatively, EVs were incubated with the pH 11.2 elution buffer for 5 min, neutralized to physiological pH and subsequently incubated with PK for 5 min (11.2*). Bar chart shows average degradation of all four proteins based on band intensity analysis.

Data from EV elution (Figure 1B and Supplementary Figure 3A) and EV permeability assays (Figure 1C and Supplementary Figure 3B) revealed that EV elution and permeability were dramatically affected by the type of elution protocol. We observed only mild differences in elution efficiency between EVs captured by antibodies against CD9 and CD81, indicating a shared release mechanism. In addition, responses were similar between both types of EVs. The overall pattern of EV elution corresponded to that of EV permeability. However, interestingly, some conditions showed moderate to high EV elution with only minimal EV permeability (e.g. compare run 11 and run 13 in Figures 1B and 1C). To identify optimal conditions to obtain both high EV elution and low EV permeability, response surface modelling was applied to the obtained datasets on CD9^+^ EV elution, CD81^+^ EV elution and EV permeability. In this process, a statistical model is constructed from the data, allowing identification of factors that significantly affect each response and allowing prediction of theoretically optimal elution conditions.

None of the outcome parameters for HEK293T EVs was significantly affected by the TEA concentration in the applied elution buffers. However, the pH of the elution buffer significantly affected elution of both CD9^+^ and CD81^+^ EV (higher pH > better elution). Whereas pH was the only significant factor driving elution for CD9^+^ EVs (model fit the data with R^2^ = 0.91 and p < 0.0001, Figure 1D), CD81^+^ EV elution was also slightly but significantly affected by incubation time, which was therefore included in the resulting model (R^2^ = 0.94, p < 0.0001, Figure 1E). No significant interaction between these parameters was detected. In contrast, both buffer pH and incubation time significantly and synergistically affected EV permeability (Figure 1F). The same pH and incubation time-dependence was observed for MDA-MB-231 EVs (Supplementary Figures 3C-E). This indicated that in order to maximize EV elution and minimize EV damage, exposure of the beads to a high pH for a short timeframe was likely optimal.

The generated models predicted that EV elution and EV permeability would start to increase at pH levels above 11. To validate these models and to identify an optimal tradeoff between EV elution and EV damage, we formulated a set of elution buffers with fixed pH levels between 11.0 and 11.5. In these buffers, we replaced TEA with disodium hydrogen phosphate, as TEA was found to be dispensable for both EV elution and EV permeability. In addition, TEA affected cell viability at high concentrations (> 20 mM), but offered poor pH stability at lower concentrations. The new elution buffers were incubated with magnetic beads coated with HEK293T EVs using antibodies against CD9 and CD81 and elution was performed for 5 min (the shortest incubation time tested in our library screen). Indeed, as predicted by the models, elution efficiency gradually increased from pH 11.1 until pH 11.5 (Figure 1G). The same behavior was observed when EVs were captured by antibodies against CD63 or a combination of Myc-tagged PS-binding proteins and anti-Myc antibodies (Figure 1G and Supplementary Figure 1C), confirming that elution occurred independent of the used antibody. The same buffers were tested in the proteinase K-based EV permeability assay for 5 min (Figure 1H). Here, a similar pH-dependent induction of EV permeability was observed.

However, a ‘burst’ effect was detected, where EV permeability remained low at pH ≤ 11.2, but suddenly increased at pH ≥ 11.3. These elution and permeability patterns were also observed for MDA-MB-231 EVs (Supplementary Figures 3F-G). Based on these observations, we selected the elution buffer with pH 11.2 as the lead candidate for further experiments. As our data showed that prolonged exposure of EVs to high pH would cause EV damage, an acidic Tris-based neutralization buffer (pH 4.1) was formulated and volumetrically titrated to neutralize the pH 11.2 buffer to physiological pH after EV elution. To investigate whether the minimal perturbation of the EV membrane observed at pH 11.2 was reversible, EVs were first exposed to the pH 11.2 buffer for 5 min, then mixed with the neutralization buffer and finally incubated with proteinase K. Indeed, intraluminal proteins remained completely protected from degradation (Figure 1H, last lane). Finally, we investigated whether the elution protocol could be universally applicable to other EV sources. We isolated EVs from HeLa cells and performed bead capture using antibodies against CD9, CD63, CD81 and using PS-binding proteins combined with anti-Myc antibodies. A similar pH-dependent elution of EVs from the beads was observed as for HEK293T EVs (Supplementary Figure 4A). EV permeability was found to follow the same pH-dependent pattern, with complete restoration of membrane integrity after the elution buffer was neutralized (Supplementary Figure 4B).

Together, these data showed that using this systematically optimized protocol, which from here onwards we refer to as ‘EV-Elute,’ EVs could be eluted from protein G-coated magnetic beads independent of antibody and EV type, with only minimal and completely reversible damage to the EV membrane.

### Bead separation allows enrichment of intact EV subpopulations with distinct proteomes

After optimizing the elution buffer, we investigated the biophysical properties of the eluted EV subpopulations. Whole HEK293T EV isolates, EV subpopulations enriched based on expression of CD9, CD63, CD81 and PS and their corresponding bead supernatants (i.e. the fraction of EVs that had not bound the beads during EV capture) were analyzed by western blotting for a range of EV-associated proteins (Figure 2A). Data showed that each tetraspanin was captured efficiently by its corresponding antibody, as shown by their near-complete removal in bead supernatants. In addition, release of each tetraspanin using EV-Elute was moderately effective with recoveries of 40-80% of their initial input levels, corresponding with previous data (Figure 1G). EV marker patterns differed among subpopulations. Interestingly, CD9^+^ and CD81^+^ EVs showed a highly similar repertoire of analyzed proteins: CD9, CD81, syntenin, TSG-101 and Alix were highly abundant in these EVs, whereas flotillin-1 and CD63 were less abundant. In contrast, CD63^+^ EVs contained less CD9, CD81, syntenin and Alix than CD9^+^ and CD81^+^ EVs, but more flotillin-1 and CD63. In concordance with our previous data (Figure 1G), PS^+^ EVs were generally obtained at higher recovery rates compared to tetraspanin-enriched subpopulations (compare PD lanes in Figure 2A with whole EV isolate), potentially due to the two-component capturing step which could be easier to disrupt using EV-Elute.

**Figure 2:**
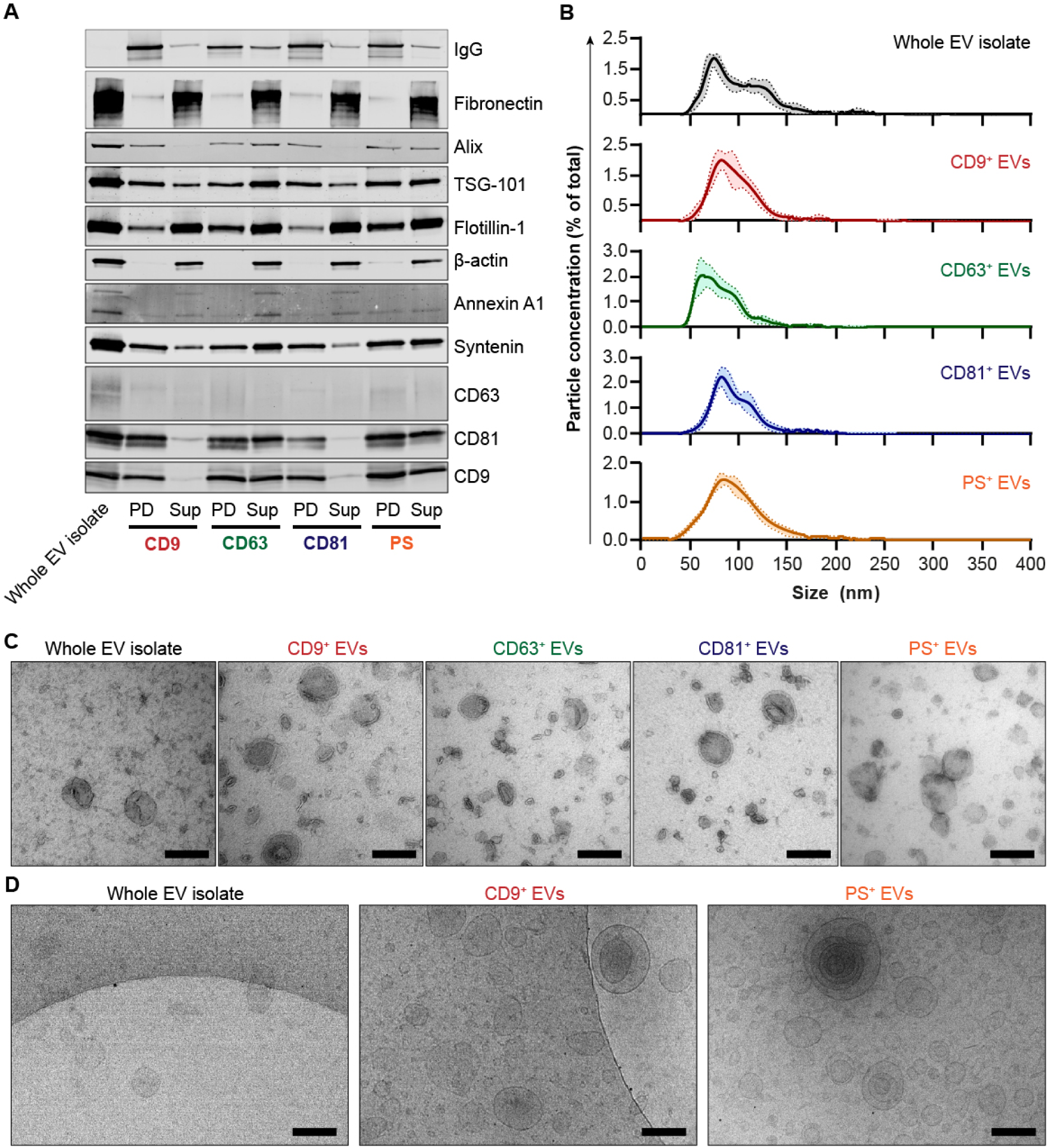
EV subpopulations isolated by EV-Elute show distinct protein profiles and morphology. **A:** Western blot analysis of common EV proteins in HEK293T EVs separated into surface-marker defined subpopulations using EV-Elute. The first lane (Whole EV isolate) shows EVs used as input for capture on magnetic beads using antibodies against CD9, CD63, CD81 and PS. Non-captured supernatant from beads was loaded in ‘Sup’ lanes, and EV subpopulations recovered with EV-Elute were loaded in ‘PD’ lanes. **B:** Representative NTA size distribution patterns of HEK293T EVs and their subpopulations. Lines show mean ± SD of 5 measurements in a single sample. **C, D:** Representative TEM (C) and Cryo-TEM (D) images of whole EV isolates and isolated EV subpopulations from HEK293T EVs. Scale bars represent 200 nm.

Interestingly, PS^+^ EVs showed an intermediate abundance of most proteins compared to CD9^+^/CD81^+^ and CD63^+^ EVs, but contained higher levels of Annexin A1. Strikingly, some proteins which were abundantly detected in whole EV isolates, such as fibronectin and β-actin, could not be efficiently captured on beads using any of the antibodies. These likely reflect contaminants in whole EV isolates, which migrate with EVs in size exclusion chromatography, but are not tightly associated with EVs.

The similarities between CD9^+^ and CD81^+^ EVs, the distinct protein profile of CD63^+^ EVs and ‘intermediate’ protein profile of PS^+^ EVs was not unique to HEK293T EVs, but was also observed for HeLa EVs (Supplementary Figure 4C).

These findings were corroborated by imaging of tetraspanins on HEK293T EV subpopulations using dSTORM imaging (Supplementary Figure 5). HEK293T EVs showed high overall abundance of CD9 and CD81, and both markers were enriched by either anti-CD9 or anti-CD81 pulldown. In contrast, CD63^+^ EVs showed lower abundance of both CD9 and CD81. PS^+^ EVs showed an intermediate phenotype with high abundance of CD63 and CD81, but lower levels of CD9.

For all EV subpopulations, a concomitant release of capturing antibodies was detected on western blot (top row in Figure 2A), indicating that the binding between bead-bound protein G and antibodies was, to some extent, disrupted by EV-Elute.

In concordance with their protein profiles, NTA revealed a highly similar particle size distribution between CD9^+^ and CD81^+^ EVs, with mode sizes of 85.3 ± 7.7 and 83.0 ± 4.4 nm, respectively (Figure 2B). In contrast, CD63^+^ EVs were smaller with a mode size of 69.1 ± 7.8 nm. PS^+^ EVs typically spanned a slightly wider range of diameters compared to the other populations with a mode size of 86.7 ± 5.9 nm. Interestingly, particle concentrations in eluted EV isolates pointed to particle recovery rates of 5-10% compared to whole EV isolates. This apparent discrepancy with previous data is likely due to the presence of non-EV contaminants in whole EV isolates, which are not captured by the beads.

Next, we analyzed EV subpopulations and their parent whole EV isolates using TEM (Figure 2C). EVs showed a commonly observed ‘cup-shaped’ morphology in all samples. However, whole EV isolates typically showed high numbers of electron-dense contaminants compared to isolated EV subpopulations, where these contaminants were almost entirely absent. These could correspond to co-isolated extracellular matrix proteins or cellular debris, in accordance with our western blotting and NTA data (Figures 2A-B). Sizes of the EV subpopulations matched those measured with NTA, where CD9^+^ and CD81^+^ EVs showed sizes similar to EVs in whole EV isolates, CD63^+^ EVs spanned smaller diameters, and PS^+^ EVs contained larger particles. Similar size distribution patterns, EV morphology and presence/absence of co-isolated contaminants was observed for HeLa EVs (Supplementary Figure 4D-E)

As traditional TEM does not show particles in their native, hydrated state, we also visualized whole EV isolates and EV subpopulations using cryo-TEM (Figure 2D). These analyses corroborated previous findings and showed that eluted EV subpopulations contained intact lipid membranes.

### EV-Elute rapidly disturbs binding of Protein G to EV-capturing antibodies

We sought to investigate the mechanism by which EVs were released from the beads. Our models showed that EV release was not dependent on incubation time in a timeframe of 5-30 min, yet EV damage progressively increased over time. However, these prediction models do not allow extrapolation beyond the original DoE design space. Hence, to investigate the release kinetics of EV elution using EV-Elute, EVs captured on beads were exposed to the elution buffer for 1 until 5 min and elution was measured using flow cytometry (Figure 3A). Interestingly, EV elution occurred almost instantaneous for all four tested capture antibodies. When elution buffer was immediately removed from beads (‘0 min’ in Figure 3A), 80% of the maximal elution efficiency was achieved for anti-tetraspanin antibodies, and elution was complete after 1 min of incubation. For PS^+^ EVs, plateau elution was instantaneously achieved. Due to technical limitations, we could not analyze EV permeability during these timeframes, but these observations led us to reduce EV elution times to 1 min for further experiments.

**Figure 3:**
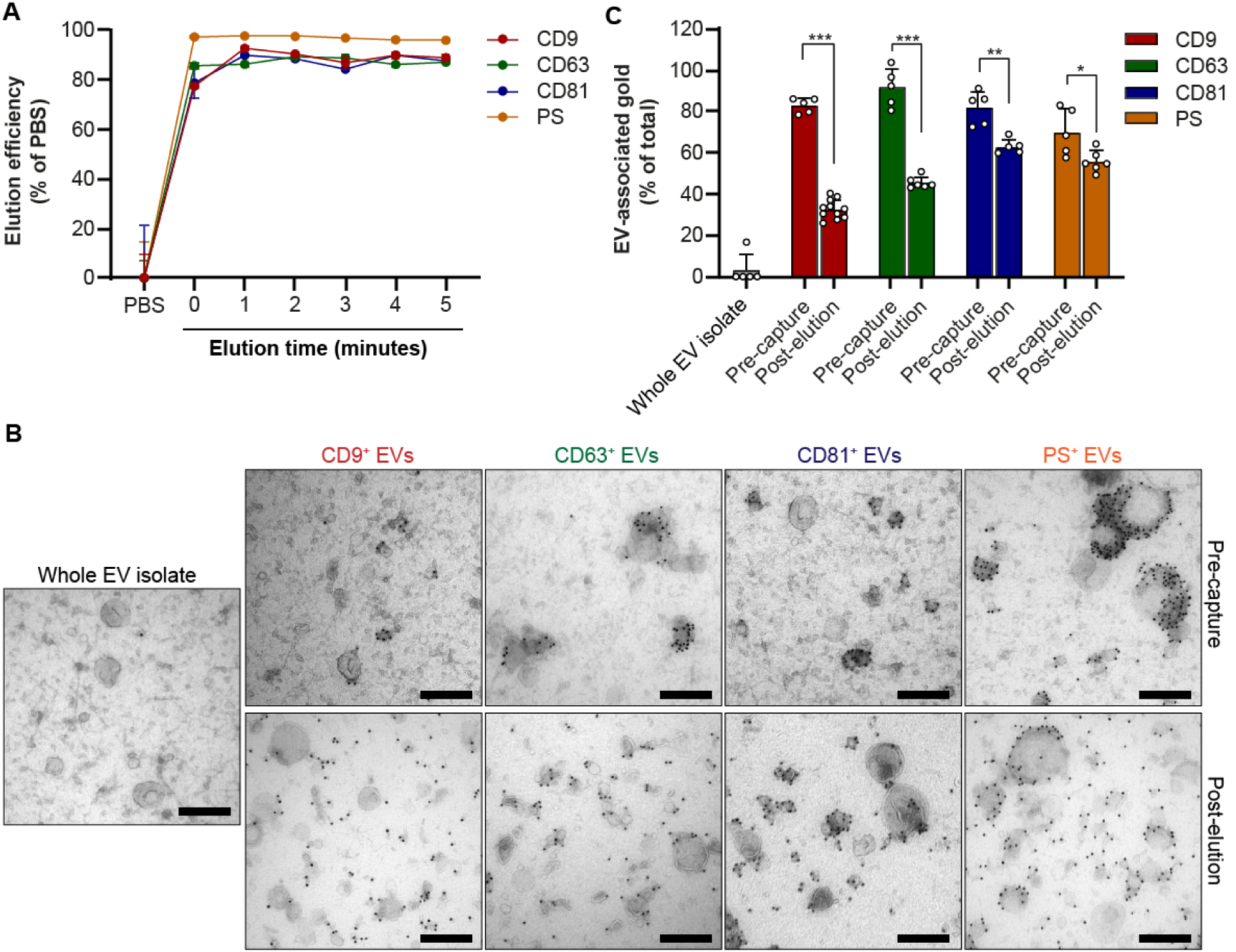
EV-Elute induces a rapid disruption of Protein G-antibody binding. **A:** HEK293T EVs were captured on Protein G-coated magnetic beads using antibodies against CD9, CD63, CD81 and PS. Beads were exposed to EV-Elute for 0-5 min, stained with PKH67 and EV elution was analyzed using flow cytometry. In 0-minute incubation, elution buffer was added to beads and immediately removed. **B:** Representative TEM pictures of whole HEK293T EV isolate mixed with capturing antibodies against CD9, CD63, CD81 and PS (’Pre-capture’) and corresponding released EV subpopulations after applying EV-Elute (Post-elution’). EM grids were immunolabelled with rabbit-anti-mouse and protein A-gold (10 nm). Scale bars represent 200 nm. **C:** Semi-automated quantification of EV-membrane associated gold in pictures shown in B. Bars show mean ± SD of 5-10 pictures. Statistical differences were determined using unpaired student’s t test; * indicates p < 0.05, ** p<0.01, and *** p<0.0001.

Western blot analysis previously revealed that capture antibodies co-released with EVs from the Protein G-beads when EV-Elute was applied. As these antibodies could interfere with downstream assays, we investigated whether the antibodies remained associated with the EV surface. To this end, we performed immunogold labelling of capturing antibodies on TEM grids coated with whole EV isolates, EVs mixed with capturing antibodies prior to bead capture and eluted EV subpopulations (Figure 3B). As expected, no gold was detected in whole EV isolates, showing that antibody staining was specific. In EV isolates mixed with capturing antibodies prior to bead capture, gold was predominantly located on the membranes of specific EV subpopulations, whereas other EVs remained completely unstained. We could not identify clear phenotypical differences between stained and unstained EVs in the same samples. However, in EV samples mixed with anti-CD63 and anti-CD81 antibodies, gold tended to localize more to EVs with smaller diameters, whereas PS-binding proteins stained predominantly larger EVs. Strikingly, in eluted EV subpopulations, a more disperse gold localization pattern was observed, which differed per antibody. In CD9^+^ and CD63^+^ EVs, roughly 50% of the gold showed clear EV membrane localization, whereas EV membrane gold localization was more pronounced for CD81^+^ and PS^+^ EVs. We observed similar patterns when these antibodies were visualized in isolated HeLa EV subpopulations (Supplementary Figure 6A).

These findings were in line with semi-automated image analysis (Figure 3C), which revealed that antibody membrane localization significantly decreased during EV elution, albeit to varying degrees depending on the antibody. Finally, we questioned whether the eluted antibodies were still functional. To this end, we exposed eluted EVs to fresh Protein G beads and analyzed their capacity to re-isolate EVs using western blotting (Supplementary Figure 6B). It was observed that all capturing antibodies could almost completely be recaptured onto the beads, suggesting that Fc-domains were still intact. At the same time, in line with our gold localization experiments, EV markers were only partially recaptured.

Taken together, these studies showed that EV-Elute causes a near-instantaneous disruption of the binding between Protein G (covalently bound to magnetic beads) and Fc domains of antibodies, resulting in release of bead-captured EVs. In this process, antibody Fc domains remain functional, but antigen-binding domains are disrupted and, depending on the antibody, released from the EV surface to variable extents.

### EV subpopulations show cell uptake and specificity similar to whole EV isolates

We next investigated whether EV subpopulations obtained using EV-Elute still retained their biological functionality. Therefore, we first interrogated whether the EV subpopulations could still be taken up by specific cells in an environment reminiscent of human blood vessels. EVs from HEK293T cells were labelled with the lipophilic dye Memglow 590 and separated into the aforementioned subpopulations using EV-Elute. In parallel, peripheral blood mononuclear cells (PBMCs) were isolated from human blood and cultured on top of a layer of HMEC-1 endothelial cells (Figure 4A). We normalized EV doses based on Memglow fluorescence, as we considered traditional protein– or particle-based normalization unsuitable for samples with vastly different purities (i.e. whole EV isolates and isolated EV subpopulations). EVs were added to PBMC-endothelial cell co-cultures, and after 4h uptake in the various immune cell subtypes was analyzed using multiplexed flow cytometry (gating strategy shown in Supplementary Figure 7A). As shown in Figure 4B, endothelial cells showed major EV uptake, and contained 80-90% of the Memglow signal (Figure 4C). Phagocytic antigen-presenting cells, including monocytes and dendritic cells, took up a major fraction of the remaining EVs. In contrast, T-cells, B-cells and NK cells hardly showed EV uptake. Some variability in absolute uptake efficiency was observed between the whole EV isolates and EV subpopulations, which likely resulted from small variations in input fluorescence and EV labelling efficiency between experiments. However, when uptake was analyzed over experiments with multiple PBMC donors, we found no significant differences in cell type specificity between EV subpopulations and whole EV isolates (Figure 4C).

**Figure 4:**
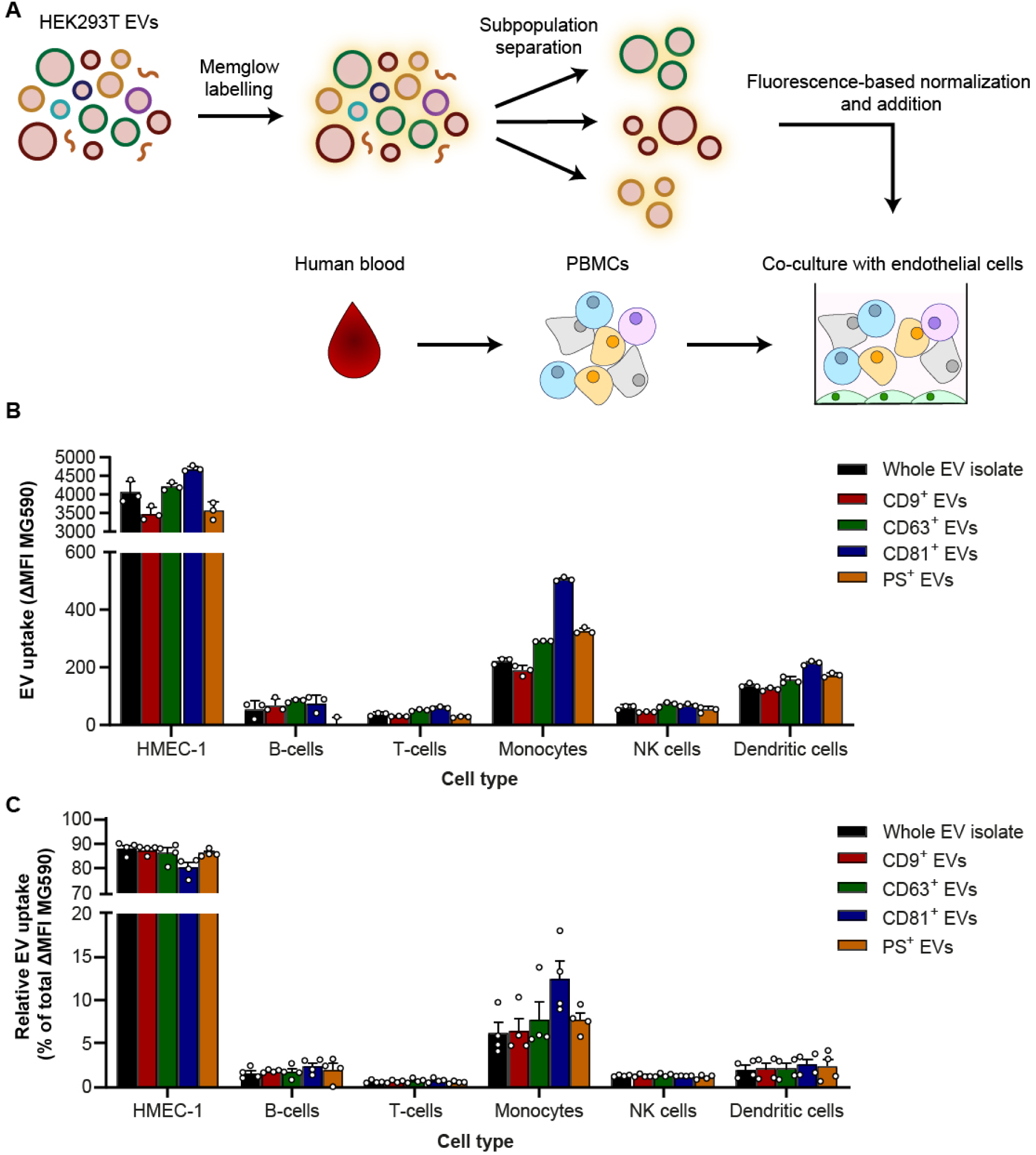
EV subpopulations are taken up efficiently and specifically by multiple cell types. **A:** Schematic illustrating the experimental setup of EV subpopulation uptake assays. EVs were labelled with Memglow 590 and added to co-cultures of human PBMCs and eGFP-expressing endothelial cells. After 4 hours, uptake by individual cell types was analyzed by multiplexed flow cytometry. **B:** Representative uptake experiment showing absolute uptake of whole EV isolates and subpopulations by co-cultures from a single PBMC donor. Uptake is expressed as the difference in mean fluorescence intensity (MFI) of Memglow 590 between EV-treated cells and PBS-treated controls (ΔMFI). Bars indicate mean ± SD from three technical replicates. **C:** Cell type specificity of whole EV isolates and EV subpopulations. Uptake of EVs by each cell type was expressed as a percentage of total EV uptake within experiments. Bars show mean ± SD of four experiments with different PBMC donors. No significant differences in cell type specificity between EV subpopulations were detected using one-way ANOVA with Tukey post-hoc test.

These data suggest that EV-Elute does not compromise the ability of EVs to be taken up by recipient cells, nor their cell type-specificity. In addition, our data indicate that despite the distinct biophysical characteristics of CD63^+^ and PS^+^ EVs compared to highly similar CD9^+^ and CD81^+^ EVs, these EV subpopulations show similar cell-type specific uptake patterns in this multicellular environment.

### Engineered therapeutic cargo delivery capacity differs between EV subpopulations

Having established that cellular uptake was not affected by EV-Elute, we investigated whether EV cargo delivery capacity remained intact. To this end, we transfected HEK293T cells with vectors driving expression of four molecules: (I) CD9 fused via a photocleavable linker[47] to the MS2 bacteriophage coat protein (MCP), which specifically binds MS2 RNA aptamers [48] (CD9-PhoCl-MCP); (II) a single guide (sg)RNA targeting a previously developed Cas9-activatable stoplight reporter[44], modified in the second loop and tetraloop with MS2 aptamers[Elsharkasy et al, manuscript in preparation]; (III) Cas9 endonuclease and (IV) glyco-protein G of the vesicular stomatitis virus (VSV-G). Together, these components drive the formation of EVs loaded with sgRNA due to its association with CD9 as well as the Cas9 protein due to its association with the sgRNA (Figure 5A). The photocleavable linker can be cleaved by irradiation with ∼400 nm UV light, releasing the Cas9/sgRNA complexes within the EV lumen and enabling their activity in recipient cells upon fusion of the EVs with endosomal membranes. This fusion process is greatly enhanced by VSV-G, allowing detectable delivery of the Cas9/sgRNA complexes with a single addition of EVs [Elsharkasy et al, manuscript in preparation]. We used this system in conjunction with a stoplight reporter construct stably expressed in HEK293T cells (HEK293T-Stoplight)[44], which consists of constitutively expressed mCherry followed by a linker region and a stop codon. The linker region is targeted by the Cas9/sgRNA ribonucleoprotein (RNP) in the EVs, and its cleavage results in permanent co-expression of eGFP due to non-homologous end joining (NHEJ)-mediated frameshift mutations (Figure 5B). Hence, cells to which Cas9/sgRNA RNP is functionally delivered by EVs express eGFP which can be visualized by fluorescence microscopy and quantified by flow cytometry.

**Figure 5:**
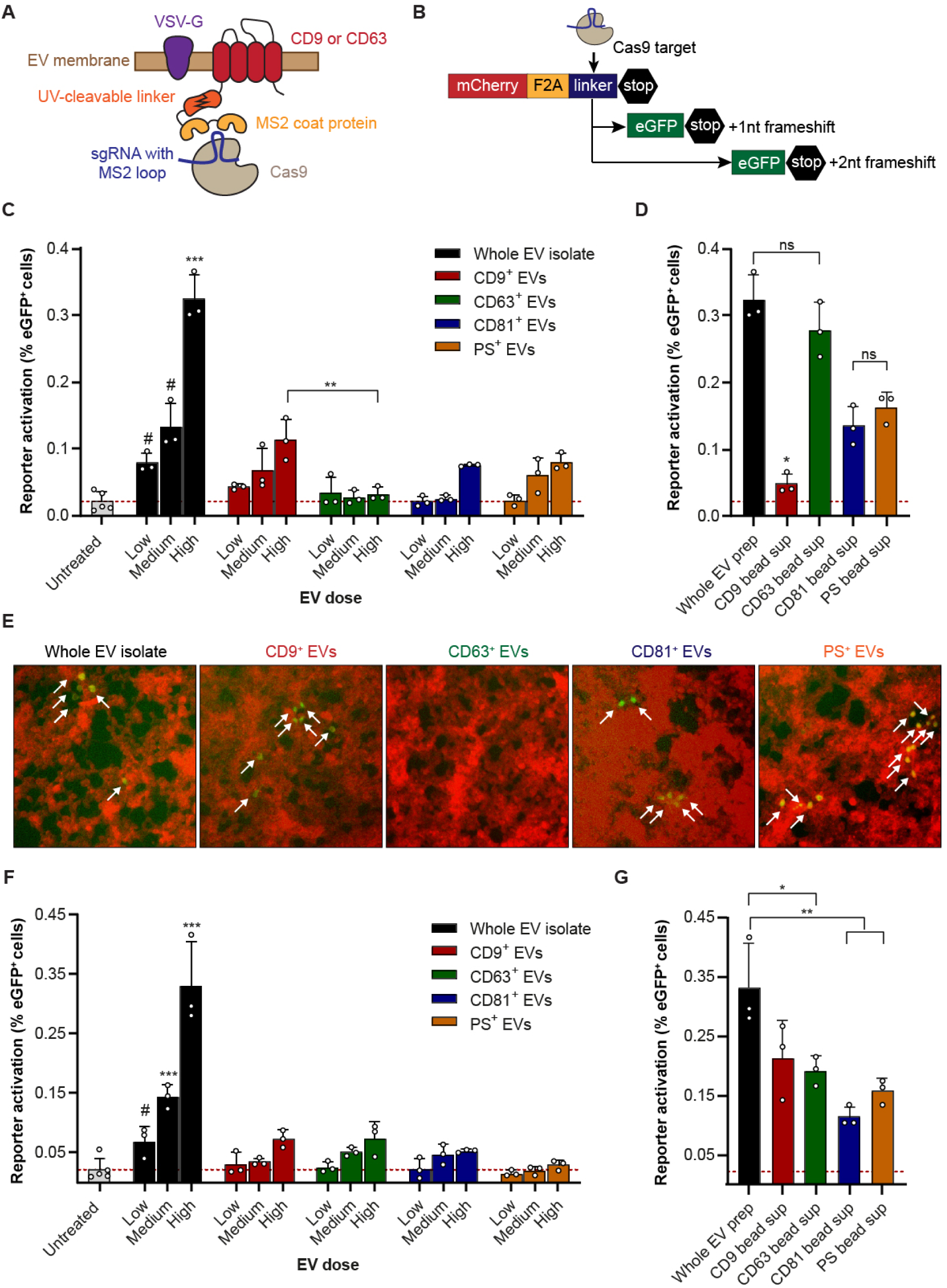
Capacity of EVs to deliver engineered cargo differs between EV subpopulations. **A:** Schematic representation of engineered EVs derived from HEK293T cells transfected with constructs encoding CD9 or CD63 fused to MS2 coat protein via UV-cleavable linkers, MS2 aptamer-modified sgRNA, Cas9 and VSV-G. **B:** Schematic representation of the Cas9/sgRNA-activatable stoplight reporter construct stably expressed in HEK293T-Stoplight cells. **C:** Activation of HEK293T-Stoplight reporter cells by escalating doses of whole isolates of engineered EVs (Cas9/sgRNA tethered to CD9) and derived surface-marker defined EV subpopulations separated by EV-Elute. Reporter activation is expressed as percentage eGFP^+^ cells of mCherry^+^ cells, measured four days after EV addition by flow cytometry. Dotted red line indicates background reporter activation. **D:** Activation of HEK293T-Stoplight cells by non-bound bead supernatants from EV subpopulations shown in C. If no ‘active’ subpopulations are captured by the beads, a reporter activation similar to that induced by whole EVs would be expected. **E:** Representative fluorescence microscopy pictures of HEK293T-Stoplight reporter cells treated with high doses of EVs analyzed in C. Activated reporter cells are indicated with white arrows. **F:** Activation of HEK293T-Stoplight reporter cells by escalating doses of whole isolates of engineered EVs (Cas9/sgRNA tethered to CD63) and derived surface-marker defined EV subpopulations separated by EV-Elute. **G:** Activation of HEK293T-Stoplight cells by non-bound bead supernatants from EV subpopulations shown in F. All graphs are representative of at least 3 replicate experiments. Corresponding EV doses were compared with one-way ANOVA with Tukey post-hoc tests. *** indicates p < 0.001 compared to all other samples unless specific comparisons are indicated, ** p < 0.01 and * p < 0.05. # indicates p < 0.05 compared to all samples except CD9^+^ EVs (ns) in C, and p < 0.05 compared to all samples except CD9^+^ and CD63^+^ EVs (ns) in F. ns = not significant.

EVs containing Cas9/sgRNA tethered to CD9 were isolated, labelled with Memglow 590 and separated into subpopulations using EV-Elute. After fluorescence-assisted concentration normalization, EVs were UV-illuminated for 20 min to release the Cas9/sgRNA complexes in the EVs, and added to HEK293T-Stoplight cells. As expected, of the selected subpopulations, CD9^+^ EVs showed highest reporter cell activation (Figure 5C and 5E). CD81^+^ EVs delivered Cas9/sgRNA to a comparable extent, in line with our previously observed similarities between CD9^+^ and CD81^+^ EVs. In addition, PS^+^ EVs, which typically showed moderate levels of native CD9 (Figure 2A), successfully activated the reporter cells. Strikingly however, despite our previous observation that CD63^+^ EVs also natively contain CD9, CD63^+^ EVs failed to induce reporter activation. These data were in line with supernatants of the magnetic bead separation (Figure 5D), which showed activation patterns of the reporter cells opposite to their corresponding EV subpopulations (CD63 > PS > CD81 > CD9). This suggests that the functional activity of the EVs could be captured from EV isolates depending on the targeted surface epitope, and could be released in the same ratios from the beads using EV-Elute. Unexpectedly however, whole EV isolates added at the same fluorescence dose typically showed 2-3– fold higher reporter activation than the best performing CD9^+^ subpopulation. We speculated that the remaining antibodies on the EVs or the elution protocol could affect EV functionality in this assay. Therefore, whole EV isolates were incubated with the same quantities of capturing antibodies as used for bead-capture and exposed to the same EV-Elute protocol. Strikingly, these controls showed equal activation of the reporter cells (Supplementary Figure 8A), indicating that the presence of capturing antibodies and/or buffers used in EV-Elute did not interfere with EV functionality. We investigated whether EV sgRNA content and protein composition could explain our findings using RT-qPCR and western blotting, respectively (Supplementary Figures 8B-C). Here, we observed that CD9^+^ EVs contained 3.5-fold more sgRNA than whole EV isolates and other subpopulations at the same fluorescence dose. In addition, all subpopulations contained CD9-PhoCl-MS2, Cas9 and VSV-G, but expression levels of neither protein could be correlated to functional activity.

As CD63^+^ EVs showed no activity in the previous experiments, yet contained all required theoretical components to activate reporter cells, we questioned whether CD63^+^ EVs were intrinsically less capable to deliver cargo in this setup. Therefore, HEK293T cells were transfected with all abovementioned constructs, but CD9-PhoCl-MS2 was replaced with CD63-PhoCl-MS2 (Figure 5A). As such, these cells secrete EVs in which the sgRNA/Cas9 RNP complexes were relatively enriched in CD63^+^ subpopulations compared to other subpopulations. Indeed, when sgRNA content was analyzed by RT-qPCR, a 4.5-fold sgRNA enrichment in CD63^+^ EVs was observed as compared to similar fluorescence doses of whole EV isolates and other subpopulations (Supplementary Figure 8D). In addition, all EV subtypes contained VSV-G and CD63-PhoCl-MCP (Supplementary Figure 8E).

However, these patterns were not corroborated with reporter activation, as CD63^+^ EVs induced similar (low) levels of eGFP expression as compared to CD81^+^ EVs and CD9^+^ EVs (Figures 5F-G). In contrast, PS^+^ EVs failed to induce eGFP expression in this setup. Again, whole EV isolates activated reporter cells 3-5-fold better than derived subpopulations at equal fluorescence doses. We once more confirmed that this was not due to inhibitory effects of antibodies bound to the EVs or the EV-Elute protocol (Supplementary Figure 8F).

Taken together, these assays showed that the EV-Elute platform could be used to compare cargo delivery potency of individual, surface-marker defined EV subpopulations.

## Discussion

In this study, we used DoE methodology to rapidly and cost-effectively optimize parameters for elution of functional EVs from Protein G-coated magnetic beads. This strategy required only 15 conditions to be tested in two different assays and allowed us to quickly identify critical factors for this elution process through statistical modelling. These models subsequently aided in the final design of the EV-Elute platform. This systematic DoE approach has a number of advantages over conventional one-factor-at-a-time screening, as (I) typically fewer experimental conditions need to be tested, (II) interactions between two factors can be detected, as we observed for EV permeability, which exponentially increased by both pH and incubation time, and (III) optimal conditions can be predicted after a single screening round. This methodology is therefore often applied by commercial enterprises to optimize industrial processes, and is now increasingly applied to optimize biological processes as well. For example, we and others employed DoE to optimize and predict therapeutic responses of mRNA-containing nanoparticles *in vivo* [49, 50]. In addition, DoE (also termed Quality-by-Design (QbD)) was recently highlighted in a “MassivEVs” workshop of the International Society for Extracellular Vesicles (ISEV) as an approach to accelerate pharmaceutical production of EVs[51], and already applied to optimize culture conditions in 3D cultures of mesenchymal stem cells to improve EV yield and enzymatic activity[52]. The current study broadens the applicability of DoE to development of novel tools to study EV biology, and advocates its further use in EV research.

Importantly, in order to set up a successful DoE screen, prior knowledge on the system to optimize is required. In our case, we performed prescreening with several candidate elution buffers to investigate which elution buffers could be compatible with EV functionality and which parameters could be optimized. In addition, it is essential to rationally select upper and lower boundaries for each parameter to be optimized. Too broad design spaces result in low accuracy for predicting optimal parameters and potentially missing subtle effects of each parameter, whereas too small design spaces may lead to optimal conditions outside the design space, which then requires rescreening. In our screen, we found that pH was the most critical factor for both EV elution and EV permeability. Most variability in these outcome parameters was found in the pH 11-11.5 range, whereas our screen covered pH 10 to 11.5. Despite our models accurately predicting that pH > 11 was optimal, a small rescreen at pH 11-11.5 was needed to finetune the elution process. This could have been avoided if the initial screen had been performed at a smaller pH range.

The EV-Elute platform was found to be universally applicable to EVs from three different cell sources, HEK293T, MDA-MB-231 and HeLa cells. In addition, we showed proof-of-concept of elution of four different capturing antibodies, targeting common EV surface markers. These antibodies varied in IgG subtype (anti-CD9: IgG2b; anti-CD81: IgG2a; anti-CD63/anti-Myc: IgG1), yet showed comparable elution efficiencies. Our data indicated that this universality is due to disruption of the bond between Protein G and the antibody during the elution process, rather than that between the antibody and epitope. This results in elution of EVs to which antibodies are still bound. The extent to which these antibodies remained EV-bound after elution varied per antibody, as would be expected given that antibody-epitope interactions are unique and complex. These remaining antibodies could affect downstream analyses. For example, cellular binding and uptake of antibody-bound EV subpopulations could be biased towards cell types expressing Fc receptors (FcRs), which are found throughout the immune system and several endothelial cell types[53]. This could potentially be avoided using antibodies derived from different species than those used in downstream assays. The capturing antibodies used in this study were of mouse origin, which can bind human FcRs in a subtype-specific manner [54]. Although we did not detect significant differences between cell-type specificity of EV subtypes and whole EV isolates (not containing antibodies) in our uptake studies, we observed a trend towards stronger uptake of CD81^+^ EVs by monocytes compared to other EV subpopulations. This may be explained by a stronger affinity of monocyte FcRs for mouse IgG2a (bound to CD81^+^ EVs) compared to IgG2b and IgG1 subtypes associated with other EV subpopulations [54]. Such biases could be ameliorated by carefully selecting capturing antibodies with species and subtypes compatible with intended downstream assays.

In addition, the potential inhibitory effect of EV-bound antibodies on EV functionality should be taken into account using appropriate controls. In Cas9/sgRNA delivery assays we approached this by exposing EVs to the same antibodies, reagents and incubation times as EV subpopulations, but omitting the magnetic beads. We strongly recommend that a similar approach is taken by future users of the EV-Elute platform. In addition, it is advisable to titrate the appropriate ratio of antibodies-to-EVs used for capture. We used approximately 12 antibodies per particle (for CD9/CD81 antibodies) as determined by NTA, but increased this to 120 antibodies per particle for CD63 antibodies to improve EV capture. These antibodies were incubated with EVs prior to mixing with magnetic beads, as we and others[55] found that this improved capture efficiency compared to immobilizing antibodies on beads first (data not shown). We subsequently incubated with an excess of Protein G-coupled beads to maximize antibody/EV capture. This process is scalable to accommodate a variety of downstream functional assays. However, despite efficient capture of designated surface markers, only ∼50% of those markers could be released using our optimized elution protocol. Our models revealed that more efficient release is theoretically possible, but we deliberately restricted release in order to maintain EV integrity. As a result, EV yields from the beads were typically limited and required EV production at a larger scale. In addition, it is conceivable that EVs with high densities of given surface markers elute less efficiently compared to their low-density counterparts. This may be solved by performing antibody-to-EV titrations, using the lowest quantity of capturing antibody possible to facilitate EV capture, but to avoid blocking associated functionality and to allow EV release. In addition, following recommendations from this study, principles from the EV-Elute platform could be incorporated in other platforms to improve yields of EV subpopulations. For example, microfluidic systems which have been previously used to characterize the molecular content of EV subpopulations (e.g. refs [56–58] and reviewed in [59, 60]) could be adapted to accommodate studies to functionality of EV subpopulations and improve EV yields compared to the magnetic bead-based platform discussed here.

Using EV-Elute, we compared the functionality of EV subpopulations containing actively loaded Cas9/sgRNA RNP complexes. In these studies, striking similarities were found between CD9^+^ and CD81^+^ EVs, which corroborated their overlapping proteomic signature we observed by western blotting. Together these data showed that in HEK293T cell-derived EVs, CD9 and CD81 are mostly located on the same particles, which appear distinct from CD63^+^ EVs. This is in line with previous immunoprecipitation-based studies[26] and typical observations that CD63 is dominantly present in EVs in multivesicular bodies[61, 62], whereas CD9 and CD81 show stronger plasma-membrane localization[63–65]. Hence, when we tethered Cas9/sgRNA complexes to CD9, both CD9^+^ and CD81^+^ EVs showed equal capacity to deliver this cargo, underlining their shared biological origin and providing proof-of-concept that EV-Elute allows functional comparison between EV subpopulations. In addition, we here demonstrate for the first time that this cargo delivery capacity extended to PS^+^ EVs, which could be due to a shared origin in the plasma membrane. PS-binding proteins are often identified in larger EVs and apoptotic bodies[66–68], but also small EVs reminiscent of CD63^+^ exosomes have been reported to contain PS [69, 70]. PS is generally known as an ‘eat-me’ signal that promotes clearance and degradation of apoptotic cells and EVs by macrophages[17, 71–73], but our data suggests that despite this potentially unfavorable phenotype for drug delivery purposes, PS^+^ EVs can still contribute to functional cargo delivery. In fact, it has previously been shown that hepatitis A-virus containing EVs fuse with endosomal membranes through PS-receptors[20], advocating a role of PS beyond waste disposal signal.

Interestingly, these experiments also showed that when tethering Cas9/sgRNA to CD9, CD63^+^ EVs could not functionally deliver this cargo despite containing equal quantities of sgRNA (and likely associated Cas9) compared to their CD81^+^ and PS^+^ counterparts. We hypothesized that in this experimental setup, CD63^+^ EVs were intrinsically unable to deliver engineered cargo. However, when we tethered Cas9/sgRNA to CD63, functional Cas9/sgRNA RNP delivery by CD63^+^ EVs could be observed. Interestingly, CD9^+^ and CD81^+^ EVs showed comparable cargo delivery as CD63^+^ EV, albeit while containing 4-fold lower quantities of sgRNA. We speculate that this is due to a shifted localization of CD63 from multivesicular bodies to the plasma membrane upon overexpression of CD63-PhoCl-MCP, which has previously been reported for other CD63 fusion proteins[74]. Hereby, the engineered cargo would become incorporated at the plasma membrane (together with CD9 and CD81), enforcing its functional delivery by plasma membrane-derived EVs rather than exosomes.

Hence, together these data suggest that plasma membrane-derived EVs (ie. CD9^+^, CD81^+^ and to some extent PS^+^ EVs) are better able to deliver engineered cargo than EVs derived from multivesicular bodies (ie. CD63^+^ EVs). This this likely due to other factors than the tetraspanins themselves, as it was recently demonstrated that CD63 and CD9 are not directly involved in EV mediated cargo delivery [75]. Of note, a potential confounding factor in our experiments was the presence of the viral fusogen VSV-G on the EVs. This protein was required to enable activation of our reporter system with a single dose of EVs, but could also significantly alter endogenous EV biogenesis pathways. In addition, VSV-G is trafficked predominantly to the plasma membrane[76, 77], although we detected considerable levels of VSV-G in CD63^+^ EVs, in line with previous co-localization studies[78]. This could have inadvertently created a bias towards stronger activity of plasma membrane-derived EVs. Nevertheless, our findings merit further investigation to functionality and delivery potency of EV subpopulations.

Intriguingly, we observed that in Cas9/sgRNA delivery assays, whole EV isolates showed severalfold higher delivery efficiencies than their derived EV subpopulations. We excluded the possibility that this was due to differences in cargo content, presence of capturing antibodies on the EVs or damage inflicted on the EVs in the elution protocol. A potential explanation for this phenomenon could be the higher purity of EV subpopulations compared to whole EV isolates. After capture on the beads, the subpopulations are washed with buffers containing low concentrations of non-ionic surfactant (Tween20) in order to remove non-specifically bound EVs. This washing step could remove part of the protein corona surrounding the bead-bound EVs, which has previously been shown to contribute to EV functionality[79, 80]. In addition, as shown by our western blot analyses, some proteins in whole EV isolates are not captured when targeting canonical EV markers. These proteins, such as the extracellular matrix component fibronectin, could be loosely surrounding EVs and potentiate their binding, uptake and potentially processing by cells[81–83]. The EV-Elute platform could aid to further elucidate the role of such proteins in future studies.

In conclusion, we present EV-Elute as a novel tool to separate and functionally compare surface marker-defined EV subpopulations. The platform relies on the transient disruption of protein G, which has affinity for IgGs derived from most species used in research applications, including mouse, rabbit, rat and human. At the same time, we showed that EV-Elute did not compromise integrity of EVs from three different sources. These features demonstrate the universal compatibility of EV-Elute with a broad range of capturing antibodies and EV sources. Thereby, the platform opens new opportunities for comparing functionality of surface-marker defined EV subpopulations beyond the canonical EV markers studied here. In addition, the platform can be used to validate functional involvement of specific proteins on EVs in a broad range of pathophysiological processes. Moreover, EV-Elute can be used to establish functional relationships between diagnostic EV surface markers and disease prognosis. Finally, EV-Elute could be used to improve therapeutic activity of EVs by enriching EV subpopulations loaded with specific cargoes.

## Acknowledgements

This work is supported by a VENI grant (no. 17296) in the Applied and Engineering Sciences Domain of the Dutch Research Council (NWO); OGJ was supported by a VENI grant in the Science Domain of the Dutch Research Council (NWO) (grant no. VI.Veni.192.174).

