## Supplementary information for "EV-Elute: a universal platform for enrichment of functional surface marker-defined extracellular vesicle subpopulations"

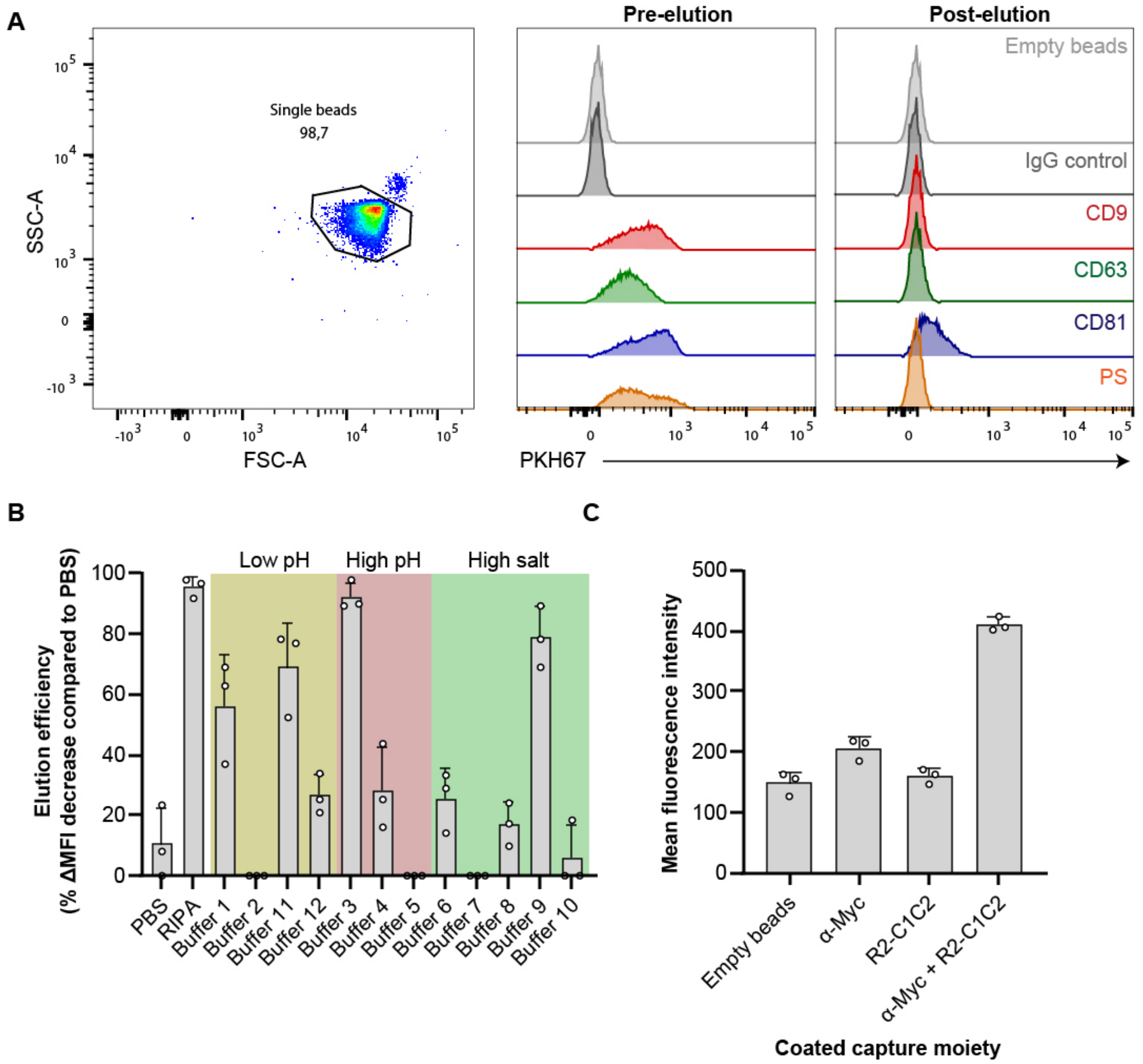


**Supplementary Figure 1: Approach to study EV elution from Protein G-coated magnetic beads using flow cytometry. A:** Gating strategy of single Protein G-Dynabeads (left) and representative histograms of beads containing HEK293T EVs captured using antibodies against CD9, CD63, CD81, IgG controls or PS binding proteins combined with anti-Myc antibodies. Beads were stained with PKH67 before (middle) and after (right) exposure to EV-Elute for 1 min. **B:** MDA-MB-231 EVs were captured on Protein G-Dynabeads using anti-CD9 antibodies and exposed to a first exploratory library of elution buffers* for 10 min. Beads were stained by PKH67 and analyzed by flow cytometry. **C:** Two-component dependency of bead capture of $PS^+$ EVs. HEK293T EVs were captured on Protein G-Dynabeads using antibodies against Myc-tag, PS-binding R2-C1C2 proteins (ref 42), or a combination of both, stained with PKH67 and analyzed by flow cytometry. Bar charts show mean ± SD of three technical replicates.

*RIPA: 1% NP-40, 0.5% octyl glucoside, 5 mM ethylenediaminetetraacetic acid (EDTA), 0.1% SDS, 50 mM Tris, 150 mM NaCl; Buffer 1: 100 mM glycine-HCl, pH 2.5; Buffer 2: 100 mM citric acid, pH 3; Buffer 3: 100 mM TEA, pH 11.5; Buffer 4: 150 mM ammonium hydroxide, pH 10.5; Buffer 5: 100 mM glycine-NaOH, pH 10; Buffer 6: 5M lithium chloride; Buffer 7: 3.5M potassium chloride; Buffer 8: 2.5M potassium iodide; Buffer 9: 3M sodium thiocyanate; Buffer 10: 100 mM Tris-acetate, 2M NaCl, pH 7.7; Buffer 11: 200 mM acetic acid, 0.5M NaCl, pH 2.5; Buffer 12: 100 mM citric acid, 2M NaCl, pH3

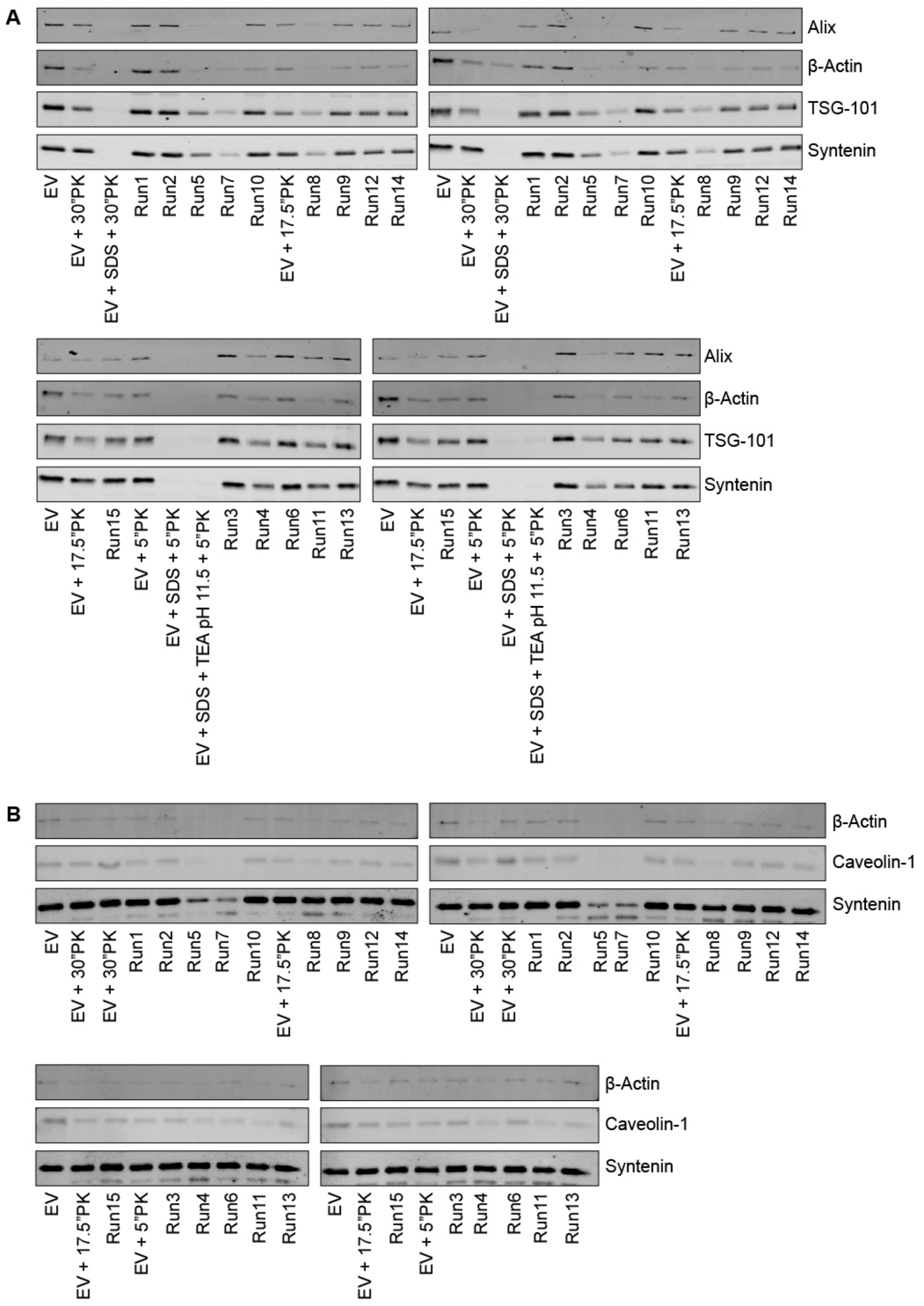


**Supplementary Figure 2: Western blots of DoE-based EV permeability assays in Figure 1C and Supplementary Figure 3B.** EVs from HEK293T (A) or MDA-MB-231 (B) were exposed to a library of elution buffers for indicated times (Supplementary Table 2) in the presence of proteinase K (PK). Abundant intraluminal EV proteins for each EV type were analyzed by western blotting. Band intensities of each protein were compared with incubation time-matched controls (either 5, 17,5 or 30 min, indicated with 5", 17.5" and 30", respectively) of EVs exposed to PK, located on the same membrane. Average degradation of the proteins is expressed in Figure 1C and Supplementary Figure 3B. All conditions were tested and blotted in duplicate.

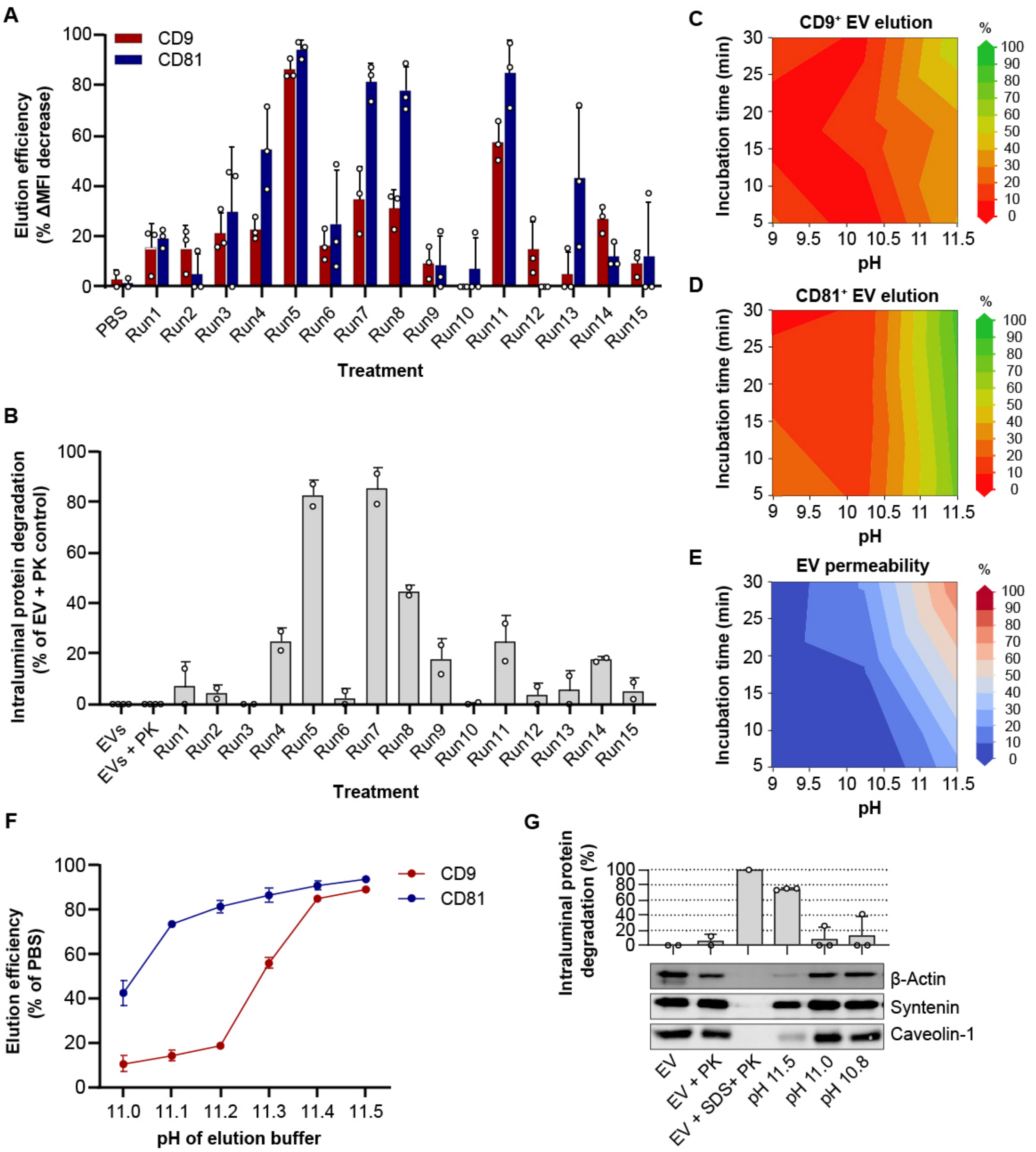


**Supplementary Figure 3: pH- and incubation time-dependence of EV elution and EV damage of MDA-MB-231 EVs. A:** MDA-MB-231 EVs were captured on Protein G-Dynabeads using antibodies against CD9 and CD81, exposed to a DoE-based library of elution conditions (Supplementary Table 2), stained with PKH67 and analyzed by flow cytometry. Bars show mean ± SD PKH67 MFI decrease of three technical replicates. **B:** MDA-MB-231 EVs were exposed the DoE-based library of elution conditions in the presence of proteinase K**.** Average degradation of intraluminal proteins β-actin, caveolin-1 and syntenin is expressed (see also Supplementary Figure 2). Bars show mean ± SD of two technical replicates. **C, D:** Prediction models of EV elution after capture with anti-CD9 (**C**) or anti-CD81 (**D**) antibodies, generated based on data in panel A. **E:** Prediction model of EV permeability generated based on data in panel B. **F, G:** Validation of EV elution and EV permeability of MDA-MB-231 EVs in the pH 11-11.5 range. **F:** EVs were captured using antibodies against CD9 or CD81 and exposed to 0.5 mM TEA-based elution buffers with pH ranging from 11.0 to 11.5 for 5 min. Beads were stained with PKH67 and EV elution efficiency was analyzed by flow cytometry. Dots show mean ± SD of three technical replicates. **G:.** EVs were exposed to 0.5 mM TEA-based elution buffer with pH 11.5, 11.0 or 10.8 for 5 min in the presence of proteinase K. Intraluminal protein degradation was analyzed by western blotting (representative blot shown on bottom) and quantified in three technical replicates (top).

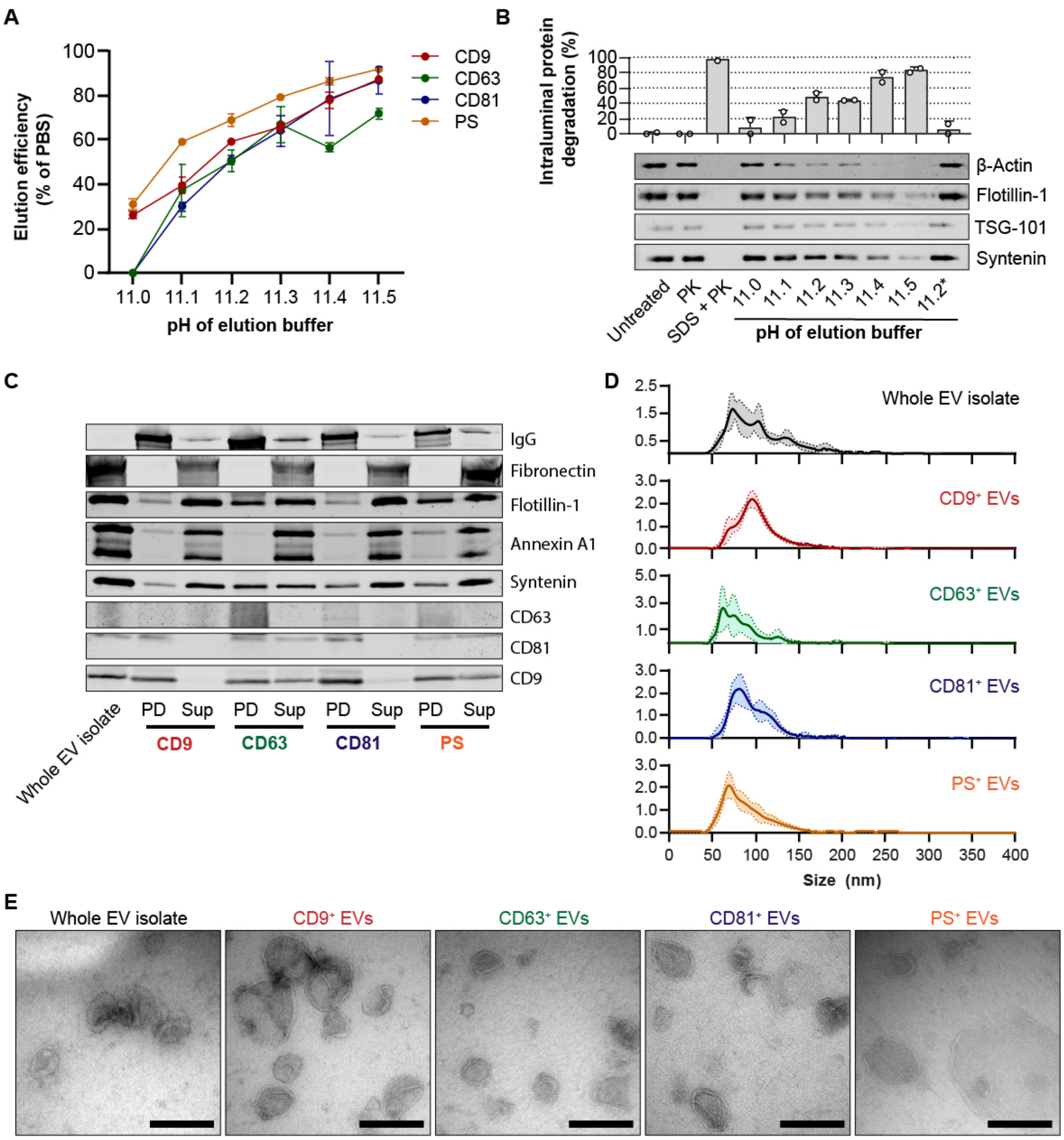


**Supplementary Figure 4: EV subpopulation separation using EV-Elute is also applicable to HeLa EVs. A:** HeLa EVs were captured on Protein G-Dynabeads using antibodies against CD9, CD63, CD81 or PS-binding proteins/anti-Myc antibodies, and exposed to disodium hydrogen phosphate-based elution buffers with pH ranging from 11.0 to 11.5 for 5 min. Beads were stained with PKH67 and EV elution efficiency was analyzed by flow cytometry. Dots show mean ± SD from 3 technical replicates. **B:** HeLa EVs were exposed to disodium hydrogen phosphate-based elution buffers with pH ranging from 11.0 to 11.5 for 5 min in presence of proteinase K (PK), after which pH was neutralized. Degradation of abundant intraluminal proteins β-actin, flotillin-1, TSG-101 and syntenin was analyzed by western blotting (bottom) and quantified (top) in two technical replicates. To test reversibility of EV damage inflicted by the pH 11.2 buffer, EVs were exposed to the pH 11.2 buffer for 5 min, pH was neutralized and subsequently EVs were exposed to PK for 5 min (last lane, 11.2*). **C:** Protein composition of HeLa EV subpopulations obtained using EV-Elute, analyzed by western blotting. Eluted EV subpopulations after pulldown are indicated with 'PD', and bead supernatants are indicated with 'Sup'. **D:** Representative size-distributions of HeLa EV subpopulations as measured using NTA. Data is shown as mean ± SD from 5 replicate measurements **E:** Representative TEM pictures of HeLa EVs (whole EV isolate) and derived EV subpopulations. Scale bars represent 200 nm.

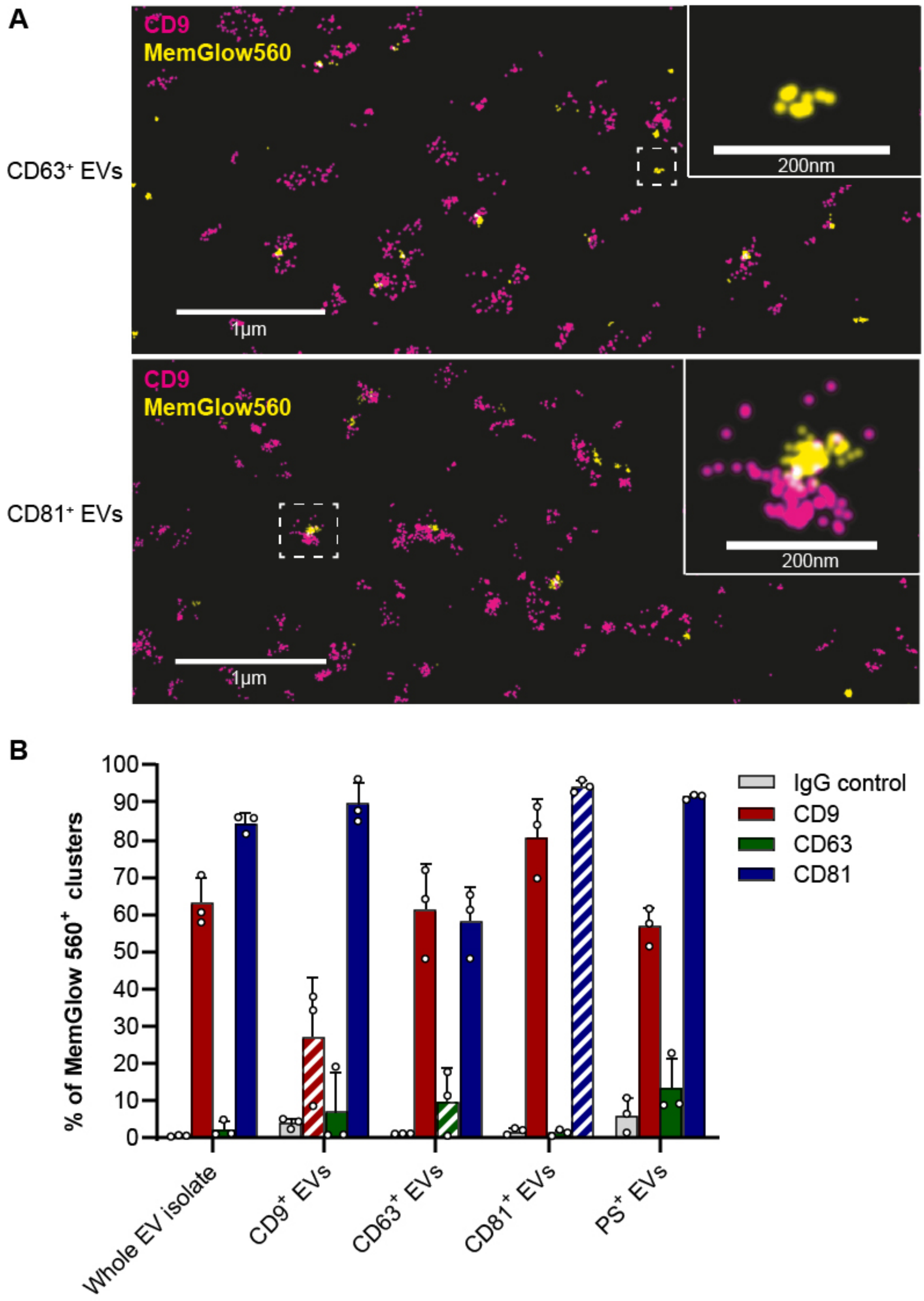


**Supplementary Figure 5: Co-localization analysis of tetraspanins on HEK293T EV subpopulations using dSTORM imaging.** **A:** HEK293T EVs were labeled with Memglow 560 (yellow), separated into subpopulations using EV-Elute and coated on coverslips. Coverslips were stained with Alexa Fluor 647-conjugated antibodies against CD9, CD63 or CD81 (magenta) and imaged using dSTORM imaging. Representative overlay pictures of CD63⁺ and CD81⁺ EVs stained with anti-CD9 antibodies are shown. Cluster analysis was performed to quantify the number of Memglow 560+ clusters that also stained positive for tetraspanins. Insets show representative tetraspanin-negative (top) and tetraspanin-positive (bottom) Memglow 560 clusters. **B:** Quantification of Memglow 560 clusters positive for CD9, CD63 and CD81 within the whole EV isolate or derived EV subpopulations. Striped bars indicate stainings for the same tetraspanins as enriched with EV-Elute. Co-localization values in these bars are likely underestimations of actual co-localization due to the presence of blocking capture antibodies on the EVs. Nevertheless, these data show that CD63+ EVs show distinct tetraspanin enrichment profiles compared to highly similar CD9⁺ and CD81⁺ EVs. PS⁺ EVs show intermediate enrichment of all tested tetraspanins. Bar chart shows mean ± of three technical replicates.

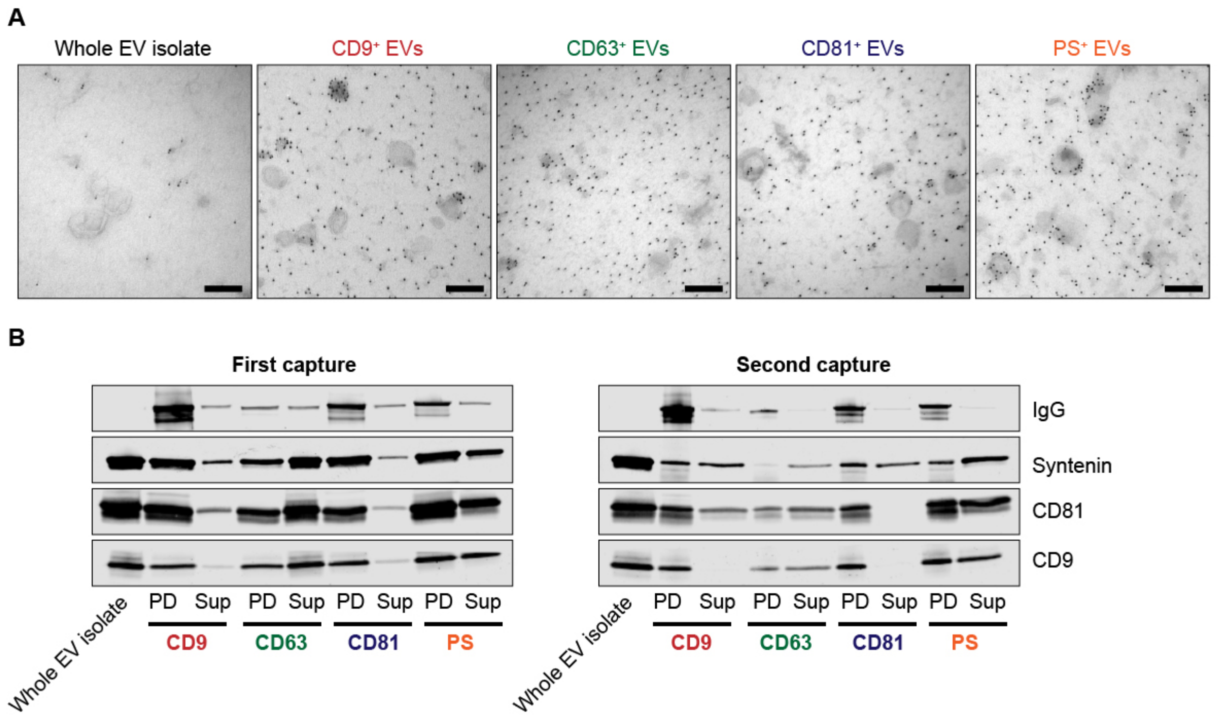


**Supplementary Figure 6: Antibodies remain partly associated with EVs after elution from magnetic beads using EV-Elute.** **A:** HeLa EVs were separated into subpopulations using EV-Elute and coated on electron microscopy grids. Capture antibodies were stained using 10 nm protein A-gold. Gold membrane localization differed per capture antibody and was similar to that observed with HEK293T EVs (Figure 3B-C). Scale bars indicate 200 nm. **B:** HEK293T EVs were captured on Protein G-Dynabeads targeting CD9, CD63, CD81 or PS and eluted using EV-Elute. EV markers were analyzed using western blotting (left panel). Eluted EVs after pulldown are indicated by 'PD', and bead supernatants are indicated with 'Sup'. Eluted EV subpopulations were pulled down again with protein G beads (without additional antibodies) and elution was performed a second time with EV-Elute (right panel). Eluted EVs and bead supernatants were compared with the same volume of whole EV isolate as in first capture, which was incubated for the same time without beads.

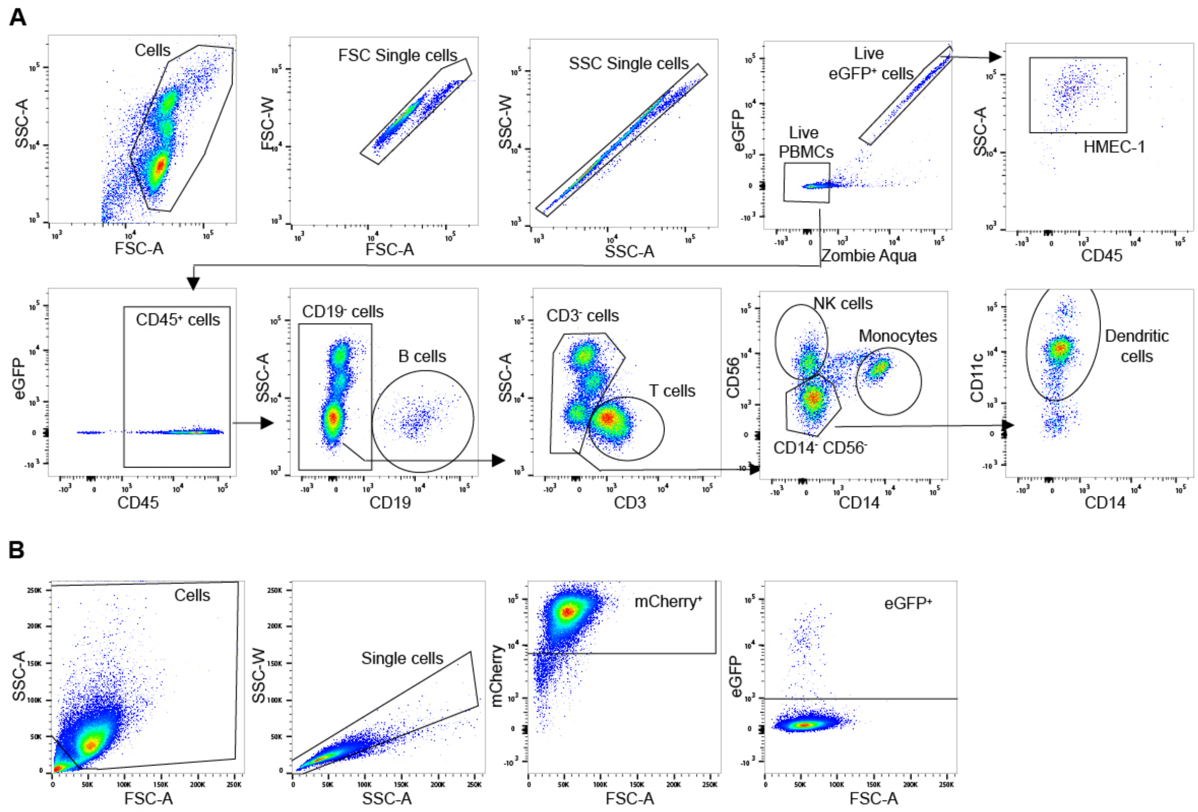


**Supplementary Figure 7: Gating strategies for PBMC-endothelial cell co-culture and HEK293T-Stoplight reporter assays.** **A:** PBMCs were co-cultured with eGFP⁺ HMEC-1 endothelial cells**.** Cells were stained with antibodies to discriminate EV uptake in endothelial cells, B-cells, T-cells, NK cells, monocytes and dendritic cells by flow cytometry. **B:** Gating strategy to quantify the percentage of mCherry⁺ HEK293T-Stoplight cells that express eGFP after successful Cas9/sgRNA delivery.

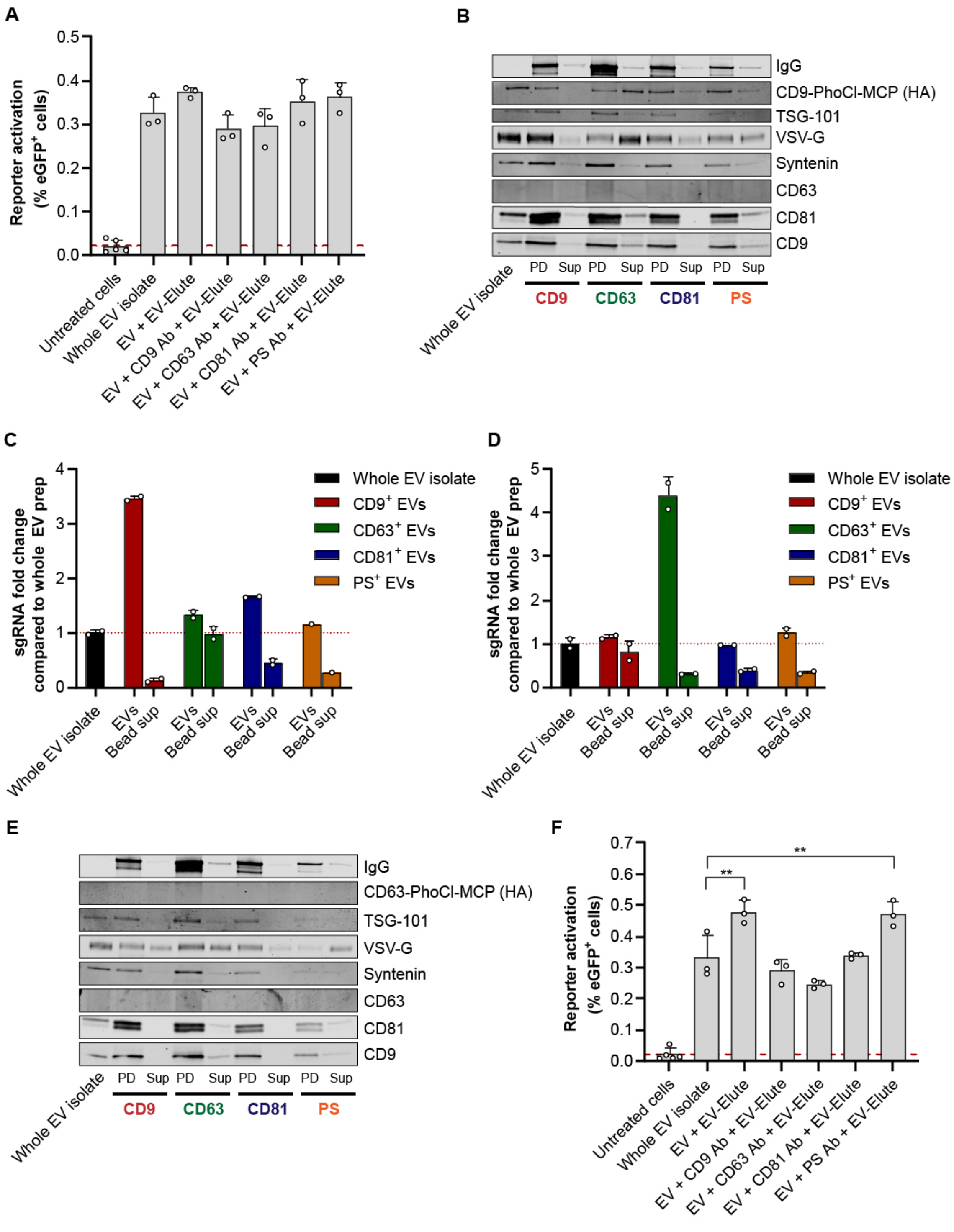


**Supplementary Figure 8: Reduced Cas9/sgRNA delivery efficiencies of EV subpopulations compared to whole EV isolates are not due to EV-Elute, presence of capturing antibodies or lack of cargo. A:** HEK293T EVs were engineered to contain VSV-G, CD9-PhoCl-MCP, MS2-tagged sgRNA and Cas9. EVs were incubated with capturing antibodies without magnetic beads and exposed to the EV-Elute protocol. After UV-illumination, EVs were added to HEK293T-Stoplight cells and eGFP expression was quantified by flow cytometry. **B:** VSV-G, CD9-PhoCl-MCP and EV markers in eluted EV subpopulations (PD) and bead supernatants (Sup) were analyzed using western blotting. EVs and bead sups were loaded in the same relative volumes as added to reporter cells in Figure 5C. **C:** sgRNA quantification in EVs and bead supernatants using RT-qPCR. The same relative volumes as added to reporter cells in Figure 5C were used as input for RNA extractions. Abundance of sgRNA in each EV subpopulation was expressed as a fold change to abundance in whole EV isolates. **D:** RT-qPCR analysis of sgRNA in HEK293T EVs engineered to contain VSV-G, CD63-PhoCl-MCP, MS2-tagged sgRNA and Cas9 and derived subpopulations. The same relative volumes as added to reporter cells in Figure 5F were used as input for RNA extractions. **E:** VSV-G, CD63-

PhoCl-MCP and EV markers in eluted EV subpopulations (PD) and bead supernatants (Sup) were analyzed using western blotting. EVs and bead sups were loaded in the same relative volumes as added to reporter cells in Figure 5F. **F:** EVs containing CD63-tethered sgRNA/Cas9 complexes were incubated with capturing antibodies without magnetic beads and exposed to the EV-Elute protocol. After UV-illumination, EVs were added to HEK293T-Stoplight cells and eGFP expression was quantified by flow cytometry **.** Statistical differences compared to whole EV isolate in panels A and F were tested using one-way ANOVA with Dunnett's post-hoc test. ** indicates $p < 0.01$.

**Supplementary Table 1: Overview of antibodies used in this study**

| **Capture antibodies** | | | | |
|---|---|---|---|---|
| **Target** | **Species** | **Clone** | **Manufacturer** | |
| CD9 | Mouse | MM2/57 | Merck | |
| CD63 | Mouse | MEM-259 | Abcam | |
| CD81 | Mouse | TS81 | Diaclone | |
| Myc-tag | Mouse | 9E10 | Home-made from 1-9E10.2 hybridoma, ATCC | |
| IgG control | Mouse | Cat. PA5-33236 | Thermo Fisher Scientific | |
| **Western blotting – primary antibodies** | | | | |
| **Target** | **Species** | **Clone** | **Manufacturer** | **Dilution** |
| Alix | Mouse | 3A9 | Thermo Fisher Scientific | 1:1000 |
| TSG-101 | Rabbit | Polyclonal, cat. ab30871 | Abcam | 1:1000 |
| B-Actin | Mouse | 8H10D10 | Cell Signaling Technology | 1:1000 |
| CD81 | Mouse | B-11 | Santa Cruz Biotechnology | 1:1000 |
| Caveolin-1 | Rabbit | EPR15554 | Abcam | 1:1000 |
| Syntenin | Mouse | OTI2H6 | Origene | 1:500 |
| Annexin A1 | Rabbit | EPR19342 | Abcam | 1:1000 |
| HA-tag | Rabbit | Polyclonal, cat. GTX115044 | GeneTex | 1:1000 |
| CD63 | Rabbit | Polyclonal, cat. A5271 | Abclonal | 1:1000 |
| CD63 | Mouse | MEM-259 | Abcam | 1:1000 |
| VSV-G | Rabbit | 8G5F11 | Kind gift from the Lyles Lab, Wake Forest University School of Medicine, USA | 1:1000 |
| CD9 | Mouse | MM2/57 | Merck | 1:1000 |
| Flotillin-1 | Rabbit | EPR6041 | Abcam | 1:1000 |
| Fibronectin | Rabbit | Polyclonal, cat. F3648 | Sigma Aldrich | 1:1000 |
| **Western blotting – secondary antibodies** | | | | |
| **Target** | **Conjugate** | **Clone** | **Manufacturer** | **Dilution** |
| Mouse IgG | Alexa Fluor 680 | Polyclonal, cat. A-21057 | Thermo Fisher Scientific | 1:7500 |
| Rabbit IgG | Alexa Fluor 680 | Polyclonal, cat. A21076 | Thermo Fisher Scientific | 1:7500 |
| Mouse IgG | IRDye800CW | Polyclonal, cat. 926-32212 | LI-COR Biosciences | 1:7500 |
| Rabbit IgG | IRDye800CW | Polyclonal, cat. 926-32211 | LI-COR Biosciences | 1:7500 |
| **Flow cytometry** | | | | |
| **Target** | **Conjugate** | **Clone** | **Manufacturer** | **Dilution** |
| CD45 | APC-Vio770 | 5B1 | Miltenyi Biotec | 1:1000 |
| CD3 | PerCP | HIT3a | BioLegend | 1:200 |
| CD19 | PE | A3-B1 | Antibodies-online | 1:100 |
| CD14 | Brilliant Violet 650 | M5E2 | BioLegend | 1:200 |
| CD56 | Brilliant Violet 786 | NCAM16.2 | BD Biosciences | 1:200 |
| CD11c | Alexa Fluor 647 | S-HCL-3 | BioLegend | 1:500 |
| Fc Block | - | Fc1 | BD Biosciences | 1:500 |
| **dSTORM imaging** | | | | |
| **Target** | **Clone** | | **Manufacturer** | |
| CD9 | HI9a | | BioLegend | |
| CD63 | MEM-259 | | Exbio | |
| CD81 | TAPA-1 | | BioLegend | |
| Mouse IgG control | 15H6 | | SouthernBiotech | |

**Supplementary Table 2: Compositions of DoE-based libraries of elution conditions**

| | HEK293T EVs | | | MDA-MB-231 EVs | | |
|---|---|---|---|---|---|---|
| | pH | Incubation time (min) | TEA concentration (mM) | pH | Incubation time (min) | TEA concentration (mM) |
| **Run1** | 10 | 30 | 20 | 9 | 30 | 20 |
| **Run2** | 10 | 30 | 0,5 | 9 | 30 | 0,5 |
| **Run3** | 10 | 5 | 20 | 9 | 5 | 20 |
| **Run4** | 11,5 | 5 | 20 | 11,5 | 5 | 20 |
| **Run5** | 11,5 | 30 | 0,5 | 11,5 | 30 | 0,5 |
| **Run6** | 10 | 5 | 0,5 | 9 | 5 | 0,5 |
| **Run7** | 11,5 | 30 | 20 | 11,5 | 30 | 20 |
| **Run8** | 11,5 | 17,5 | 10,25 | 11,5 | 17,5 | 10,25 |
| **Run9** | 10,75 | 17,5 | 10,25 | 10,25 | 17,5 | 10,25 |
| **Run10** | 10,75 | 30 | 10,25 | 10,25 | 30 | 10,25 |
| **Run11** | 11,5 | 5 | 0,5 | 11,5 | 5 | 0,5 |
| **Run12** | 10,75 | 17,5 | 0,5 | 10,25 | 17,5 | 0,5 |
| **Run13** | 10,75 | 5 | 10,25 | 10,25 | 5 | 10,25 |
| **Run14** | 10,75 | 17,5 | 20 | 10,25 | 17,5 | 20 |
| **Run15** | 10 | 17,5 | 10,25 | 9 | 17,5 | 10,25 |

**Supplementary Table 3: List of used oligonucleotides and primers for RT-qPCR amplification of sgRNA against the Cas9-activatable stoplight construct (T-sgRNA) and a spike-in non-targeting control sgRNA (NT-sgRNA)**

| Name | Sequence (5’-3’) |
|---|---|
| **Oligonucleotides** | |
| NT-sgRNA spike-in | UCUCUAUCACUGAUAGGGAGGUUUUAGAGCUAGAAAUAGCAAGUUAAAAUAAGGCUAGU CCGUUAUCAACUUGAAAAAGUGGCACCGAGUCGGUGCUUUUUU |
| **Primers (used for both RT and qPCR steps)** | |
| Reverse T-sgRNA | TGTTGGCCAAGTTGATAACGG |
| Reverse NT-sgRNA | GACTCGGTGCCACTTTTTCAA |
| Forward T-sgRNA | CAGTACTCCGCTCGAGTGTT |
| Forward NT-sgRNA | TCACTGATAGGGAGGTTTTAGAGC |
